# A logarithmic theory of visuomotor stabilization

**DOI:** 10.64898/2025.12.11.693625

**Authors:** Leonardo Demarchi

## Abstract

Although many animals rely on visual information to navigate, optic flow is inherently ambiguous as it confounds information about motion speed and object distance. As a result, the visual feedback produced by a given motor command is context-dependent and requires an appropriately adapted response. Recent experiments have investigated how the fish *Danionella cerebrum* use visual cues to stabilize their position against simulated external currents. Logarithmic sensorimotor transformations have been proposed to enable adaptive responses to perturbations while preventing delay-induced instabilities. Here, we develop the theoretical framework introduced for continuous locomotion to show how logarithmic coding naturally gives rise to this adaptive behavior. The system is modeled by a nonlinear delay differential equation, which is analyzed using dynamical systems theory. We further analyze experimental data to uncover the mechanisms underlying swimming initiation and positional drift correction. Finally, we extend our framework to intermittent locomotion, resulting in a nonlinear difference equation, and show that it still produces robust adaptive behavior. This is motivated by the literature on zebrafish, where visuomotor stabilization has been extensively studied, but burst-and-coast swimming obscures the underlying adaptation mechanism. We show that our theory can reproduce the experimental results reported for motor adaptation in zebrafish without invoking internal models. Overall, our results highlight logarithmic coding as a unifying principle for visuomotor stability across continuous and intermittent locomotor regimes.

## I. INTRODUCTION

Vision is widespread in the animal kingdom, as it provides individuals with detailed information about their surroundings, important for survival and reproduction [1]. In particular, it plays a central role in guiding locomotion and spatial navigation [2]. While navigating complex environments, animals must continuously adapt their behavior to external perturbations in order to maintain their course. This is a difficult task, as sensory inputs only give a limited and noisy view of the state of the body and the environment, and motor actions are hindered by finite temporal delays [3]. These challenges are not unique to biology, they are also faced by engineers designing robotic systems capable of operating in realworld scenarios [4]. One approach that has emerged as a common solution, and is employed by animals and robots in multiple contexts, is the implementation of forward internal models, which is to say, the ability to use the current motor commands to predict future sensory inputs [5–8]. Such models enable organisms to anticipate delays and adapt their behavior by using the measured sensory inputs to minimize prediction errors, thereby maintaining accurate internal predictions across varying contexts. Although powerful, this approach is computationally demanding, suggesting that evolution may have found simpler mechanisms to achieve robust adaptation for basic problems common across species.

An example of such a common problem is that of stabilizing one’s position against external currents while navigating through a fluid. As an animal moves relative to its environment, the visual scene it perceives changes over time. The resulting apparent motion of the surrounding objects, called optic flow, is an important visual cue used to guide locomotion [9, 10]. While terrestrial animals like humans generally rely on friction with the ground to maintain stability, flying and swimming animals must continually adjust their movements to avoid being carried away by air or water currents. In these cases, vision is crucial for remaining stable relative to the surroundings, as any displacement produces a corresponding optic flow [11, 12]. Thus, these organisms share the challenge of using visual inputs to modulate their thrust and compensate the effect of external perturbations.

In neuroscience, a common experimental paradigm is to present visual stimuli that simulate such whole-field motion of the animal. In this settings, the animals reliably respond by moving in a way that compensates for the apparent perturbation, a behavior known as the optomotor response [13, 14]. In recent years, larval zebrafish has been used extensively as a model to study this behavior [15–17], as their small size and transparency allow for functional whole-brain imaging at cellular resolution, making it possible to investigate the neural circuits involved [18, 19]. However, researchers do not agree on the underlying computations performed by this animal. Some have proposed that the fish perform motor learning using a forward internal model [20, 21], or even that they encode a memory of their location in the environment [22]. Others have proposed alternative mechanisms based on simpler feedback controllers [23–25], but these models are usually designed to explain data from individual experiments, disregarding prior findings. The difficulty mainly arises from the fact that zebrafish swim intermittently in discrete bouts, allowing them to independently adjust the intensity and timing of each bout to achieve the same average speed. As a result, the problem of stabilization through intermittent locomotion is underdetermined, making it challenging to identify the underlying sensorimotor computations.

A different animal model, the miniature fish *Danionella cerebrum* [26, 27], is emerging as an alternative to zebrafish. It remains small and transparent at the adult stage, such that brain imaging can still be performed [28], while also permitting the study of a broader range of behaviors [29–33]. Another advantage of this model is that, at the larval stage, it swims continuously [34], making it much easier to study visuomotor stabilization by simply tracking the modulations of swimming speed over time. In fact, recent experiments have shown that the apparent adaptive behavior observed during visuomotor stabilization does not result from an internal model, but rather arises from the presence of nonlinear sensorimotor transformations [35]. These transformations correspond to the logarithmic encoding of both sensory input and motor output, which are well-known features of sensory (Weber-Fechner law) and motor (Henneman’s size principle) ends of the nervous system [36, 37], suggesting that the resulting adaptive stabilization may be common to other species. By extending this model to the case of intermittent locomotion, we show that it can account for the adaptive behavior observed in zebrafish without invoking internal models or restricting the analysis to individual experimental results.

In Sec. II, we first introduce the translational optomotor response and review the literature on zebrafish by summarizing the findings of studies investigating this behavior. Then, in Sec. III, we revisit the recent experiments in danionella and analyze the mathematical framework proposed for visuomotor stabilization in the case of continuous locomotion. We also examine the initiation of swimming in response to external currents and an additional integration mechanism that prevents positional drift. Finally, in Sec. IV, we extend this framework to the case of intermittent locomotion, to study the stabilization process in zebrafish. We show that, even in this case, logarithmic coding leads to an emergent adaptation, preventing the instabilities that could arise from a delayed response to incomplete sensory information.

## II. TRANSLATIONAL OPTOMOTOR RESPONSE

### A. Quantification of optic flow

We begin by discussing in more detail what optic flow is and how to quantify it, since it is the feature of visual input that is relevant for the stabilization behavior studied in the rest of the paper. In the absence of mechanisms for depth estimation, the visual inputs available to the animal consist in the information of the distribution of light as a function of its direction with respect to the eye. When an object moves relative to the animal, the angular position of its image on the retina changes. The optic flow refers to the angular velocity of the images moving across the retina. It can be quantified as the ratio between the component of the object velocity perpendicular to the line of sight and the distance of the object from the observer [9].

Consider a fish moving forward with speed *V* at a distance *h* from a planar visual pattern (Fig. 1a). In the reference frame that moves with fish, the pattern moves backward with the same speed *V*. Different points on the pattern will move differently across the retina, because of their different distances and directions, giving rise to a specific form of translational optic flow that reflects the specific geometry of the environment. We will only consider translational movements of such a planar pattern along the fish heading direction, therefore we can characterize the intensity and direction of the optic flow with the optic flow rate *ω* = − *V/h* [24]. This is simply the ratio between the velocity of the pattern and its distance, corresponding to the maximum possible angular velocity, which would be attained by the point located directly below the fish. We take it to be positive when the pattern moves forward with respect to the fish and negative when it moves backward. Importantly, we note that changing both the speed and the distance of the pattern by the same factor results in the same optic flow rate.

**FIG. 1.**
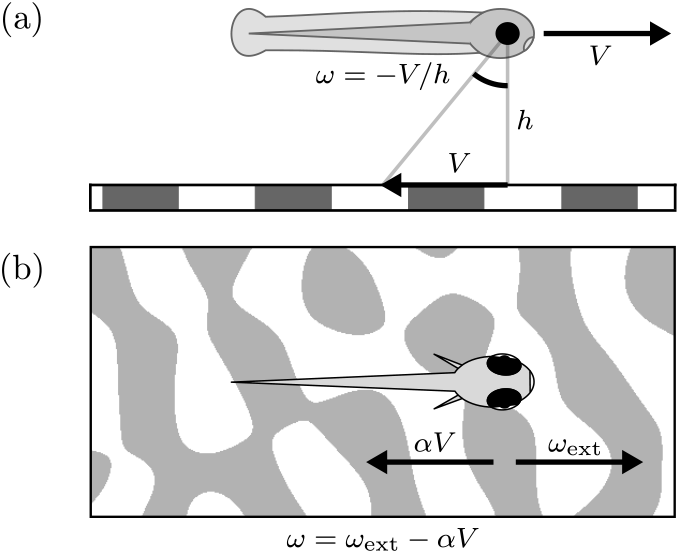
Translational optic flow during fish locomotion. (a) A fish, moving forward with speed *V* relative to a stationary visual pattern at distance *h*, experiences a backward optic flow with characteristic angular velocity *ω*. (b) The total optic flow rate *ω* results from a forward component *ω*_ext_, corresponding to the backward drift caused by a water current, and a backward component − *αV*, corresponding to the visual feedback due to forward swimming.

Now, if we externally translate the pattern it will give rise to the same optic flow as if it was the fish that was moving, but in the opposite direction. When we move the pattern forward with an external flow rate *ω*_ext_ > 0, it corresponds to a backward movement of the fish. This visual stimulus is known to elicit an optomotor response, whereby animals actively move forward to counteract the effect of this fictive backward current. When the fish moves forward with speed *V* it experiences a backward optic flow with a rate proportional to its speed − *αV*. We indicated the feedback gain *α* in place of the inverse pattern distance 1*/h* because certain experiments make it possible to manipulate the feedback and thus disentangle it from the pattern distance. The overall optic flow rate is then given by the sum of these two contributions *ω* = *ω*_ext_ − *αV* (Fig. 1b).

### B. Experiments in zebrafish larvae

There have been several studies investigating the translational optomotor response in the larval zebrafish, as its small size and transparency allow the use of imaging techniques to investigate the underlying neural circuits [15, 17]. Mainly three different experimental approaches have been used to probe this behavior. In all cases the fish is positioned in a water tank and a visual pattern is displayed below it and translated to simulate the presence of an external water current. In the first approach (freely swimming), the fish are free to move and naturally experience the visual feedback resulting from their motion with respect to the visual pattern. In the second approach (head-restrained), the fish are embedded in an agarose gel in such a way that they can move their tail but their head remains stuck in place. A camera is used to record the tail movements and estimate what would be the corresponding forward motion of the fish. The visual pattern is updated accordingly in real time to restore the visual feedback in this restrained condition. In this way, one can choose the feedback gain *α*, which determines the intensity of the visual feedback for a given estimate of the speed, which is equivalent to choosing the pattern distance *h* in the freely swimming condition. The third approach (paralyzed) is similar to the second, except that the fish are paralyzed, and their intended movements are inferred from electrode recordings of motor nerve activity in their tail.

The first study quantifying how these fish can stabilize their position on average against external currents was Ref. 38. The authors performed experiments with the freely swimming approach and varied the external flow rate across a broad range *ω*_ext_ = 0.2–8 rad/s, while the feedback gain was fixed by the distance of the visual pattern at *α* = 2 cm^−1^. They observed that the fish adapted their average speed up to ≈ 2 cm/s to match that of the external current. This adaptation resulted from an increasing bout speed *V*_b_, an increasing bout duration *T*_b_ and a decreasing interbout duration *T*_i_ as *ω*_ext_ is increased. They also noted how the reaction time *T*_r_ between the onset of the current and the initiation of swimming decreased with *ω*_ext_. Finally, they performed experiments in head-restrained conditions, but did not observe perfect stabilization, they reported significantly larger values of both *T*_b_ and *T*_i_.

Refs. 20 and 21 restrained the fish to easily manipulate the visual feedback, and studied how their behavior progressively changed after perturbations of the feedback gain. They presented fish with external currents with flow rate *ω*_ext_ = 2 rad/s and switched between three different values of the feedback gain *α*. They did not quantify whether the fish stabilized their position with respect to the visual pattern. The authors of Ref. 20 used the head-restrained approach and found that, after increasing *α*, the duration of swimming bouts *T*_b_ decreased and the interbout duration *T*_i_ increased, with these changes manifesting gradually over several seconds. The authors of Ref. 21 used the paralyzed approach and found that, after increasing *α*, the bout speed *V*_b_ decreased over a few bouts and converged to a different value. They also observed a lower rate of swimming bouts *f*_b_ = 1*/*(*T*_b_ + *T*_i_) for larger *α*. Both Refs. 20 and 21 proposed that fish maintain an internal representation of the relationship between motor command and visual feedback, updated through motor learning by comparing predicted and observed feedback.

Ref. 39 studied the reaction times of fish to external currents using the head-restrained approach. The authors argued that swimming initiation follows a Poisson process, in which the fish can start swimming at any time with a rate dependent on *ω*_ext_, as opposed to an evidence accumulation process in which the fish integrate the flow rate and start swimming when a threshold is reached.

Ref. 23 introduced a feedback controller to describe the response of the fish to perturbations, again using the head-restrained approach. The authors perturbed the visual feedback corresponding to each swimming bout in a randomized fashion, either choosing different values of the feedback gain *α*, delaying the feedback by different times, or suppressing it completely in certain phases of the bout. They found the speed profile at the beginning of the swimming bout to be independent of the perturbations, which only resulted in a modulation of the final part of the bout, after a delay of ≈ 200 ms. They modeled the swimming initiation and termination with a feedback controller and used it to fit the observed bout and interbout durations *T*_b,i_. The model does not account for changes of the swimming speed, but only for the timing of the swimming bouts based on the value of a motor drive with respect to a state-dependent threshold. The motor drive is given by the difference between the leaky integration of the net flow rate and that of the motor output, respectively modeling the integration of the sensory input and a process of motor inhibition.

Ref. 24 used three different distances of the visual pattern to vary manipulate the visual feedback in freely swimming conditions. The authors noted that the speed during bouts followed the same temporal profile once renormalized by the initial speed. They found that fish adapted their swimming speed to that of the current, but did not match it exactly, resulting in imperfect stabilization. The bout speed *V*_b_ increased with *ω*_ext_ and decreased with *α*, while the bout rate *f*_b_ increased with *ω*_ext_ and was independent of *α*. They fitted the dependence of *V*_b_ and *f*_b_ using a feedback controller built upon the models of Refs. 39 and 23. Bout initiation was modeled as a Poisson process with a rate proportional to the instantaneous forward flow rate and modulated by motor inhibition, whereas bout speed was controlled by the leaky integration of the forward flow rate.

Ref. 22 proposed that the fish maintain a memory of their location in the environment, from experiments with the paralyzed approach. The authors presented fish with brief currents in open-loop conditions (*α* = 0), directed either forward or backward, and followed, after a short pause, by a forward current in closed-loop conditions (*α* > 0). They found that fish adapted their fictive swimming in the closed-loop phase to stabilize their position, compensating for the simulated displacement in the open-loop phase. The observed trajectories are compatible with a feedback controller that keeps the integrated net flow rate close to zero.

Ref. 25 reproduced the experiments of Ref. 22 with the head-restrained approach. They varied the pauses between the two phases, finding that the swimming speed in the closed-loop phase remained significantly different for forward and backward open-loop currents, even after a pause of 16 s. They also tested a protocol in which a closed-loop current was followed by an open-loop one, finding that, for a given number of bouts in the closedloop phase, the swimming speed during the open-loop phase was independent of *α*. They argued that fish can separate the external visual flow from the feedback, only using the former to regulate their speed, which implies the use of an internal model. Finally, they performed another experiment where they showed an open-loop current with a randomly switching direction, finding that the bout-triggered average stimulus decays exponentially with a time constant of ≈ 3 s. They reproduced these results using a model where swimming bouts are generated by a Poisson process with a time-varying rate. The rate was derived from leaky integration of the external flow rate, with a larger time constant in the absence of visual flow.

It is apparent that the experimental results and theoretical interpretations are not always compatible across different papers. On the experimental side, some of the inconsistencies might be due to different experimental conditions and approaches used to study the behavior. Some papers report that the fish can stabilize their position exactly, while others suggest that the regulation is imperfect. One possible explanation for this discrepancy is the effect of light refraction. In fact, projecting the visual stimulus onto a screen positioned outside the water tank, results in a distorted image from the perspective of an observer inside the water [40]. The stimulus appears compressed within a circular window corresponding to a cone with half-angle of ≈ 49^°^. In order to simulate the effect an external current one would like to reproduce an optic flow corresponding to a backward movement of the fish. Without refraction, and with a screen that covers most of the lower visual field, a forward translation of the pattern is equivalent to a backward translation of the fish. In the presence of refraction, however, the pattern inside the circular window still moves forward, while the window itself remains stationary relative to the fish, producing a conflicting stimulus. Many of the papers we discussed here seem to have ignored this problem, which could partly explain the lack of consistency among them. On the theoretical side, one would like to have a single interpretation that can describe most of the experimental observations reported in the literature. However, it is more common to find models tailored to reproduce the data presented within the same paper, often disregarding inconsistencies with earlier results. For example, Refs. 24 and 25 suggest that the fish modulate their response only based on the forward or external flow rates, but this is inconsistent with the ability of the fish to stabilize their position across different conditions, as shown in Refs. 21, 38 and 22. In Sec. IV, we address this conflict and propose a theoretical interpretation that accounts for most of the observations in zebrafish larvae. This interpretation is based on the mechanism that was proposed for visuomotor stabilization in danionella larvae [35], which we further develop in the following section.

## III. STABILIZATION THROUGH CONTINUOUS LOCOMOTION

### A. Experiments in danionella larvae

Here, we further analyze the results of Ref. 35, where visuomotor stabilization was investigated in danionella larvae. A head-restrained approach was used, with the visual pattern projected directly on the bottom of the tank, in order to avoid distortions due to light refraction. While zebrafish larvae move through a discrete series of swimming bouts, danionella larvae can swim continuously for minutes, resulting in a distinct behavioral response [34]. The fish were presented with external currents of 30 s duration for different values of external flow rate *ω*_ext_ and feedback gain *α*. Three different experiments were performed (Fig. 2, from left to right):

**FIG. 2.**
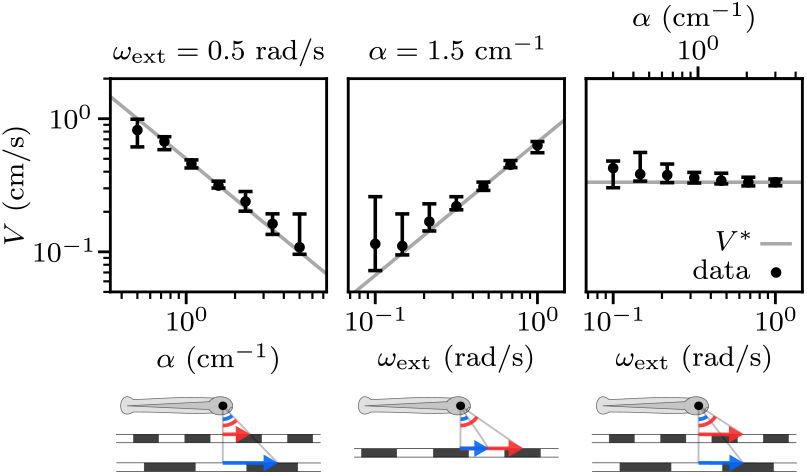
End-of-trial swimming speed *V* in response to external currents for different values of feedback gain *α* and external flow rate *ω*_ext_, from the experiments of Ref. 35. The median swimming speed was calculated for each individual fish from the periods where it was swimming in response to the current in the last 10 s of the trials (black error bars, quartiles of the distributions across *N* = (33, 31, 28) fish, from left to right). The target speed is given by *V* ^*^ = *ω*_ext_*/α* (gray lines). The schematics illustrate how parameter changes map to variations in pattern distance and current speed in the real world. The arrows indicate the current speed for two different parameter values (smaller in blue, larger in red).

- Increasing *α* with constant *ω*_ext_, corresponds to decreasing distance and current speed.
- Increasing *ω*_ext_ with constant *α* corresponds to increasing current speed with constant distance.
- Increasing *α* and *ω*_ext_ proportionally, corresponds to decreasing distance with constant current speed.

The fish adapted their swimming speed *V* to approximately swim at the target speed *V* ^*^ = *ω*_ext_*/α* for which for which the visual feedback compensates the external flow. It was shown that the observed adaptation results from a simple feedback control mechanism, combined with the fact that the nervous system encodes physical variables logarithmically. We now recapitulate how this mechanism works in the absence of noise, to make clearer what is the effect of each nonlinearity in the sensorimotor loop and how they shape the behavior of the system.

### B. Delay-induced instability in linear feedback control

It was shown that there are significant delays in the sensorimotor loop corresponding to the optomotor response, such that a change in optic flow rate at a given time only has an effect on the swimming speed after a delay *τ* ≈ 150 ms [23, 35]. Following Ref. 35, we first consider the case of a linear system where the acceleration 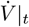 is proportional to the delayed net optic flow rate *ω*|_*t*−*τ*_, with a responsiveness *k*:

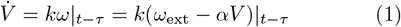

The solutions to this equation depend on the value of the dimensionless gain *µ* = *kατ*. They converge to the target speed *V* ^*^ = *ω*_ext_*/α* exponentially for *µ* < 1*/e* and through damped oscillations for 1*/e* < *µ* < *π/*2, whereas they are diverging oscillations for *µ* > *π/*2. Therefore the system becomes unstable if either responsiveness, feedback gain, or delay are increased above a critical value.

To illustrate the behavior of the system we show the solutions for three different parameter values. We consider increasing values of the external flow rate *ω*_ext_ while keeping a constant target speed *V* ^*^. Therefore, the feedback gain *α* also increases proportionally to *ω*_ext_. The increase in *α* leads to an increasing slope of the response function (Fig. 3a), eventually resulting in an unstable fixed point. The solutions showing the evolution of *V* over time (Fig. 3b) illustrate the three different dynamical regimes described above. As the characteristics of the external current are characterized by *ω*_ext_ and *α* we can summarize the behavior of the system in a two-dimensional stability diagram. Here, we equivalently chose one of the axis to be the target speed *V* ^*^ (Fig. 4a). We see that for any given *V* ^*^, the system becomes unstable as *ω*_ext_ is increased above a threshold *πV* ^*^*/*(2*kτ*), leading to diverging speed oscillations. It is important to note that this description must eventually break down, as the fish can only swim within a limited range of speed. Therefore, this model fails when *V* falls outside this range, as for example if it becomes negative.

**FIG. 3.**
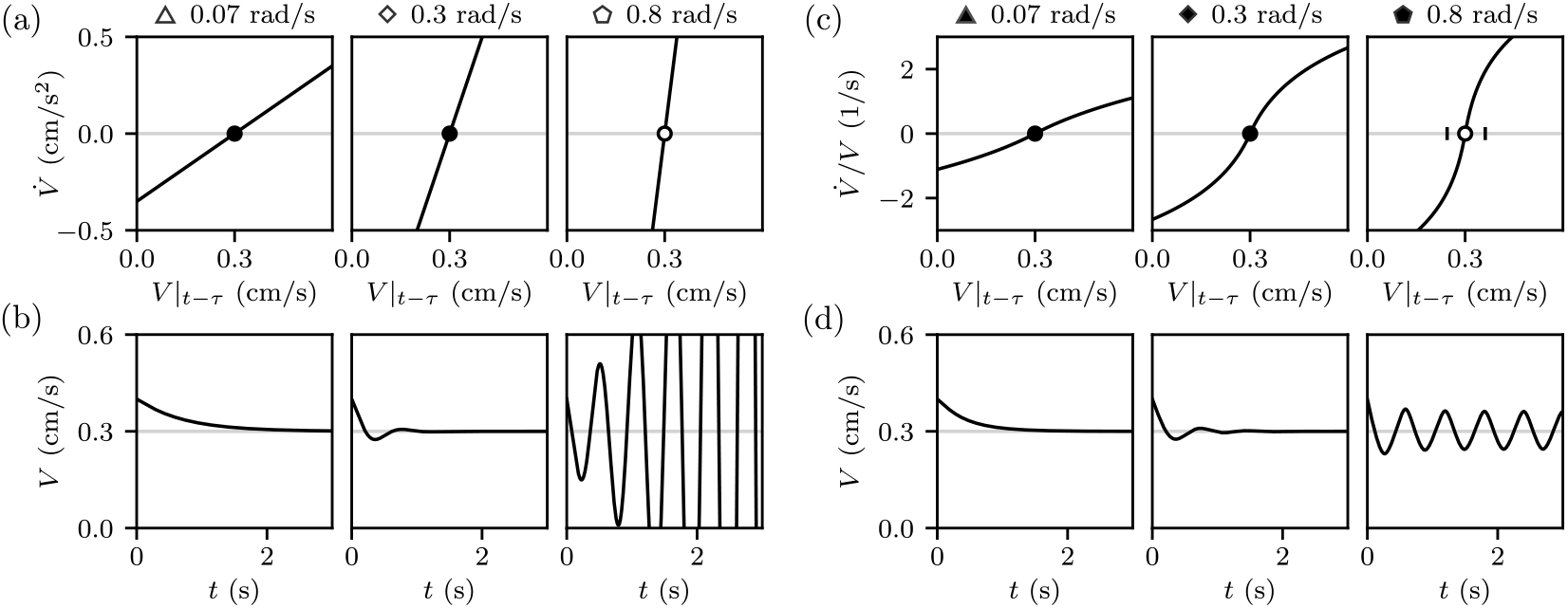
Examples of response functions and swimming speed traces for the linear and nonlinear feedback control equations. (a) Linear response functions of Eq. 1 for three choices of *ω*_ext_ (values above the subplots) for a fixed target speed *V* ^*^ = 0.2 cm/s. The black dots indicate stable fixed points and the white dot an unstable one. (b) Solutions of Eq. 1 for the corresponding parameters in (a) and an initial swimming speed *V*_0_ = 0.3 cm/s. (c) Same as (a), but for the nonlinear response functions of Eq. 3. The black vertical lines indicate the amplitude of the limit cycle. (d) Same as (b), but for the solutions of Eq. 3. The symbols mark the corresponding points in the stability diagrams of Fig. 4. The parameter values here and in the following figures, unless otherwise stated, are: *k* = 5 cm/s, *τ* = 0.15 s, *r* = 1.6 s^−1^, *ω*_c_ = 0.07 rad/s.

**FIG. 4.**
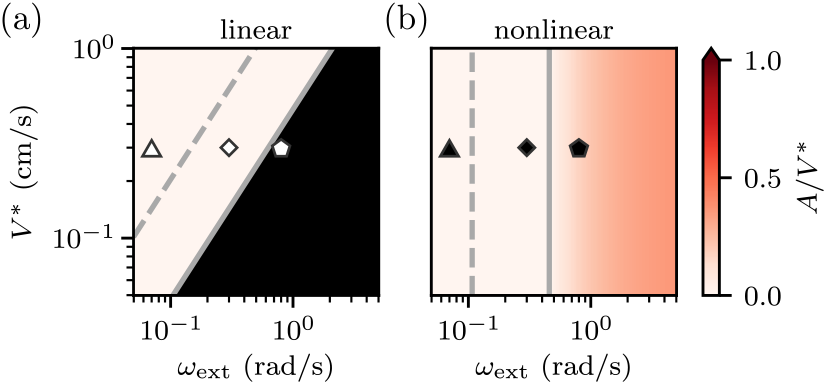
Stability diagrams for the linear and nonlinear feedback control equations. Amplitude *A* of the steady-state solution for (a) Eq. 1 and (b) Eq. 3, for different values of the target speed *V* ^*^ and the external flow rate *ω*_ext_. The region where the system is unstable is colored in black. The limit cycle amplitude was computed according to the approximation described in the text. The gray dashed lines indicate the transition to oscillatory behavior, whereas the solid ones indicate the bifurcation boundary.

Our analysis highlights that this feedback control model is problematic: for any given value of *ω*_ext_, the system becomes unstable if the target speed *V* ^*^ is too low. The fish cannot measure the value of *V* ^*^ directly from sensory inputs, as it can with *ω*_ext_. This is because *V* ^*^ depends on *α*, which can only be estimated using efference copies of motor command to disentangle exafferent and reafferent visual inputs. In principle, this could be achieved by a forward internal model which would provide an internal estimate of the feedback gain *α*. Using this information, the fish could then adapt its responsiveness *k* to reach the target quickly while also avoiding instability. However, in Ref. 35, it was shown that a different mechanism actually prevents this instability and gives rise to adaptive responsiveness without motor learning.

### C. Effect of logarithmic transformations

The behavior of the system was found to arise from the presence of nonlinear sensorimotor transformations that were not included in Eq. 1 (Fig. 5). It was shown that both the optic flow rate and the swimming speed are encoded logarithmically in the brain, in accordance with the Weber-Fechner law and Henneman’s size principle, respectively [36, 37]. This results in a logarithmic compression from the optic flow rate *ω* to a corresponding sensory drive *S* and an exponential expansion from a motor drive *M* to the corresponding swimming speed *V*. Finally, the motor drive is taken to be proportional to the integral of the sensory drive through a rate coefficient *r*, yielding the three following sensorimotor transformations:

**FIG. 5.**
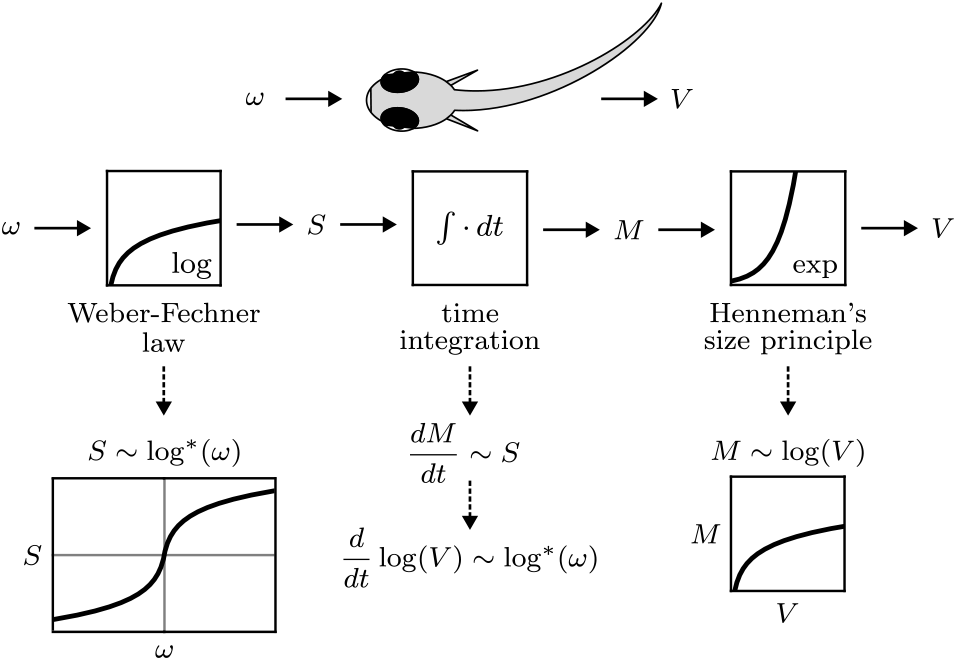
Sensorimotor transformations underlying visuomotor stabilization. The optic flow rate *ω* is logarithmically compressed at the sensory end, producing a sensory drive *S*. This signal is integrated over time to generate a motor drive *M*, which is exponentially expanded at the motor end, resulting in the swimming speed *V*.

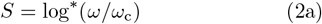

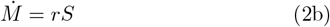

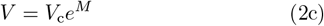

Where the parameters *ω*_c_ and *V*_c_ specify the transformation from optic flow rate to sensory drive and from motor drive to swimming speed, respectively. For the sensory drive, we considered a shifted logarithm of the form log^*^(*x*) = sgn(*x*) log(1 + |*x*|), which is continuously differentiable close to the origin to make the system easier to study analytically. We actually expect the fish to be unable to detect flow rates with a magnitude falling below a certain threshold *ω*_th_, and a more accurate transformation would be a thresholded logarithm of the form log_th_(*x*) = sgn(*x*) log(|*x* |)*H*(|*x*| − 1), with *H* being the Heaviside step function. Both functional forms lead to analogous conclusions, differing only in their behavior for small flow rates comparable with the detection threshold.

Combining Eqs. 2 and including the time delay *τ*, we obtain the following equation for the dynamics of *V* :

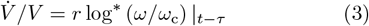

In Ref. 35, the response function of the fish was estimated, yielding the values *r* ≈ 1.6 s^−1^ and *ω*_c_ ≈ 0.07 rad/s, which we consider here. While Ref. 35 included a multiplicative noise term in Eq. 3 to reproduce the experimentally observed speed traces, here we consider only the deterministic part of the dynamics to examine the effect of the nonlinearities on the response of the system. We can still linearize Eq. 3 to study the dynamics close to the target speed *V* ^*^, recovering a linear equation of the form of Eq. 1, with the dimensionless gain now given by *µ*_0_ = *rτω*_ext_*/ω*_c_. For *µ*_0_ < *π/*2, the fixed point is stable and the solutions converge to it as in the linear case. For *µ*_0_ > *π/*2, the presence of the nonlinearities prevents the system from diverging and the solutions converge to a stable limit cycle, corresponding to regular speed oscillations around the target speed. In the limit of small oscillations it was shown that the amplitude of the swimming speed oscillations grows linearly with the bifurcation parameter as *A* ≃ 1.7(*ω*_c_*/α*)(*µ*_0_ − *π/*2). Thus, the relative oscillation amplitude *A/V* ^*^ remains bounded as *ω*_ext_ increases. We plotted the stability diagram as for the linear system, finding that the stability of the system now depends only on the external flow rate *ω*_ext_ (Fig. 4b). At *µ*_0_ = *π/*2 one observes a Hopf bifurcation, followed by limit cycle oscillations with a finite amplitude. This result holds close to the bifurcation, but as the system is driven further away from it *A/V* ^*^ is expected to grow logarithmically with *ω*_ext_, and eventually break the limit of small oscillations. On the other hand, we also expect the response to eventually saturate for large enough *ω*_ext_, similarly to how we expect it to vanish below a certain detection threshold. Such saturation could be incorporated into the functional form of the sensory drive *S*, limiting the growth of the relative oscillation amplitude and resulting in confined oscillations around the target speed even for very large values of *ω*_ext_. As in the linear case, we plotted the solutions for three increasing values of *ω*_ext_ corresponding to different dynamical regimes (Fig. 3d). The slope of the response function at the fixed point still increases with *ω*_ext_, resulting in an unstable fixed point, but now the sublinear response results in limit cycle oscillations (Fig. 3c).

We can now clearly distinguish the effects of the two nonlinear transformations. The logarithmic compression at the sensory end limits the responsiveness as the swimming speed deviates from the target in such a way that the relative amplitude of the oscillations around it remains bounded. On the other hand, the exponential expansion at the motor end results in a responsiveness that is proportional to the swimming speed itself, so that the dependence on the feedback gain is replaced by a dependence on the external flow rate. This is advantageous because, unlike *α, ω*_ext_ can be directly measured from visual inputs. Given that the fish can only detect a limited range of optic flow rates, the response magnitude *r* can be set, for example through natural selection, so that the responsiveness remains sufficiently high across this entire range.

### D. Initiation of swimming

Ref. 35 only considered how the fish continuously adapt their swimming speed to stabilize their position, not how swimming is initiated in response to the appearance of a visual flow. Here, we further analyze the same data to gain insight into this process. The termination of swimming could be included in the model by assuming that the fish stop swimming soon after their speed *V* reaches the boundaries of the range of physiologically accessible speeds [*V*_min_, *V*_max_]. In this way one can reproduce the observations that fish stop swimming after their speed reaches a maximum *V*_max_ in open-loop conditions, or after it reaches a minimum *V*_min_ after the external current is removed [35].

For the same experiments described in Sec. III A, we extracted the initial speeds *V*_0_ as the maximum speed reached by the animal in the first time *τ* after initiation of swimming (Fig. 6a) and the reaction times *T*_r_ between the appearance of the external current and the moment where the fish start swimming (Fig. 6b).

**FIG. 6.**
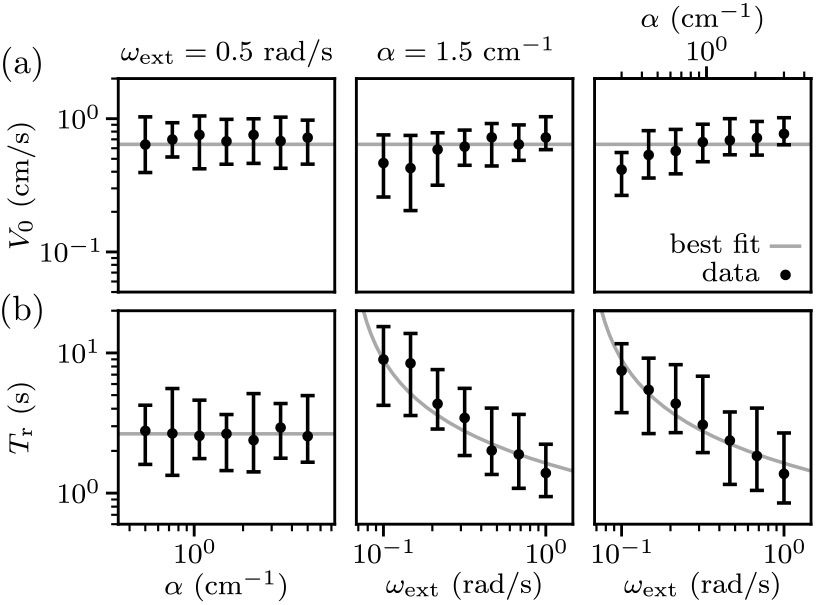
Initial speeds *V*_0_ (a) and reaction times *T*_r_ (b) in response to external currents for different values of feedback gain *α* and external flow rate *ω*_ext_, from the same experiments as in Fig. 2. Error bars denote the quartiles of the distributions (data from *N* = (33, 31, 28) fish, from left to right). The initial speed was estimated as the maximum swimming speed attained in the first 150 ms following initiation of swimming. Trials in which fish were already swimming at the onset, or never started swimming, were excluded from the distributions. *T*_r_ as function of *ω*_ext_ is well-fit by an inverse logarithmic dependence (Eq. 4), whereas all other plots are well-fit with constant (gray lines).

The initial speed *V*_0_ seems largely independent of *α* and *ω*_ext_, remaining close to the upper bound of the physiological speed range. Although we observe a slight increasing trend with *ω*_ext_, the fish generally seem to always start swimming at high speeds.

The reaction time is independent of *α*, and decreases with *ω*_ext_. Its independence from *α* is expected, as the fish cannot know the feedback gain before starting to swim. Interestingly, the width of the *T*_r_ distribution grows proportionally with *T*_r_ itself, resulting in uniform widths when plotted on a logarithmic scale. To clarify this, we plotted histograms of the logarithm of the measured values of *T*_r_ for different values of *ω*_ext_ (Fig. 7). The distributions are peaked and approximately Gaussian, suggesting that the reaction times follow lognormal distributions.

**FIG. 7.**
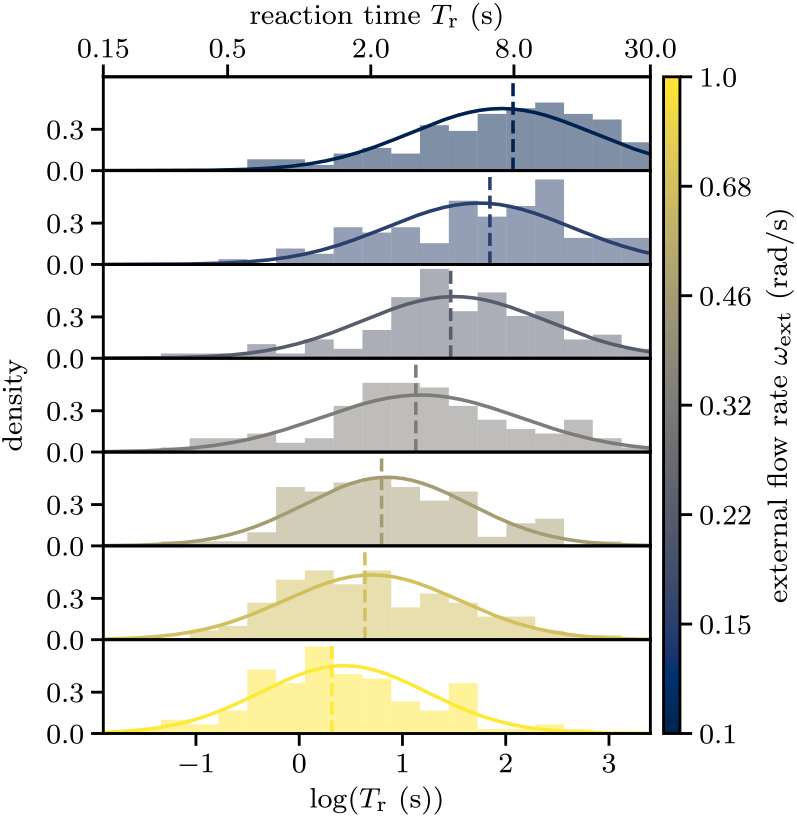
Distribution of reaction times for different values of the external flow rate. Each subplot shows the histogram of the logarithm of the reaction time *T*_r_, pooling the measurements shown in Fig. 6b for the same value of *ω*_ext_ (corresponding color bar). The distributions are well-fit by Gaussian distributions of comparable widths (solid lines). The medians of the distributions decrease with *ω*_ext_ (vertical dashed lines).

Ref. 39 proposed that larval zebrafish initiate swimming stochastically via a Poisson process, which would produce exponential distributions of reaction times. In contrast, we observe unimodal distributions with a positive peak, suggesting that swimming initiation involves an integration process. We can consider a simple deterministic accumulation model in which the sensory drive *S* is integrated over time, and the fish initiates swimming when the integrated signal Φ = *∫ S*(*t*)*dt* reaches a threshold Φ_r_. This kind of model has been proposed as a fundamental mechanism underlying decision making and reaction time processes [41]. For a constant *ω*_ext_ it would result in a reaction time *T*_r_ = Φ_r_*/S*(*ω*_ext_).

The dependence of reaction times on stimulus intensity has been traditionally fit with decreasing power laws, and interpreted either as reflecting nonlinear transformations of the stimulus or resulting from the integration mechanism itself [42]. In our simple accumulation model, *T*_r_ decreases with *ω*_ext_ with a dependence that is determined by the nonlinear compression of the sensory drive *S*. Here, we can consider the thresholded logarithm log_th_ as the optic flow rate remains positive before swimming is initiated, resulting in an inverse logarithmic dependence for the reaction times:

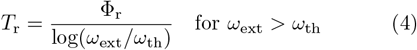

We fitted the observed dependence with this functional form and found that it fits well the data with Φ_r_ ≈ 5 s and *ω*_th_ ≈ 0.03 rad/s (Fig. 6b). In fact, the estimated value of *ω*_th_ is consistent with the one estimated in Ref. 35 by directly measuring the response function.

If we admit that the motor drive *M* starts increasing due to the presence of the external flow before initiation of swimming, then Eq. 2 predicts that the fish would start swimming at a speed 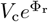, independent of *ω*_ext_ and *α*. This is also consistent with our observation that the initial speed *V*_0_ is approximately constant (Fig. 6a). To understand the observed variability in reaction times for any given value of *ω*_ext_ (Fig. 7), we hypothesize that it arises from a variability in the internal sensory drive *S*. Rather than being constant, *S* is assumed to fluctuate due to a source of multiplicative noise, such that its coefficient of variation remains constant. Then, its values will follow a lognormal distribution *S* ∼ Lognormal(log(*m*_*S*_), 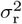), where *m*_*S*_ is the median of *S* and *σ*_r_ controls the relative magnitude of the fluctuations. This assumption seems reasonable, as neuronal firing rates, along with many other structural and functional properties of the brain, have been shown to follow lognormal statistics [43].

If *S* is lognormally distributed, then the resulting reaction time *T*_r_ = Φ_r_*/S*(*ω*_ext_) will also follow a lognormal distribution *T*_r_ ∼ Lognormal(log(Φ_r_*/m*_*S*_), 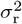). The observed fluctuations of *T*_r_ are compatible with the described mechanism, as for different *ω*_ext_ we find that a lognormal distribution provides a good fit, with approximately the same value of *σ*_r_ ≈ 0.9 (Fig. 7). In fact, experimentally measured reaction times in other contexts have been successfully fitted with the lognormal distribution, and other possible explanations for how this distribution arises have been proposed [44].

### E. Drift correction

We have seen the fish start swimming at a speed *V*_0_ close to the maximum attainable speed *V*_max_. Then, following Eq. 3 for the evolution of the speed would result in *V* approaching the target *V* ^*^ with a rate of the order of *rω*_ext_*/ω*_c_. Using the estimated values of *r* and *ω*_c_, we would therefore expect *V* to approach *V* ^*^ on a subsecond time scale in our experiments. However, this is not what is observed. Looking at the transient evolution of *V* after initiation of swimming, one finds a slow convergence occurring over several seconds (Fig. 8a-c). This becomes particularly clear when plotting the estimated sensory drive and observing how it gradually approaches zero (Fig. 8d-f).

**FIG. 8.**
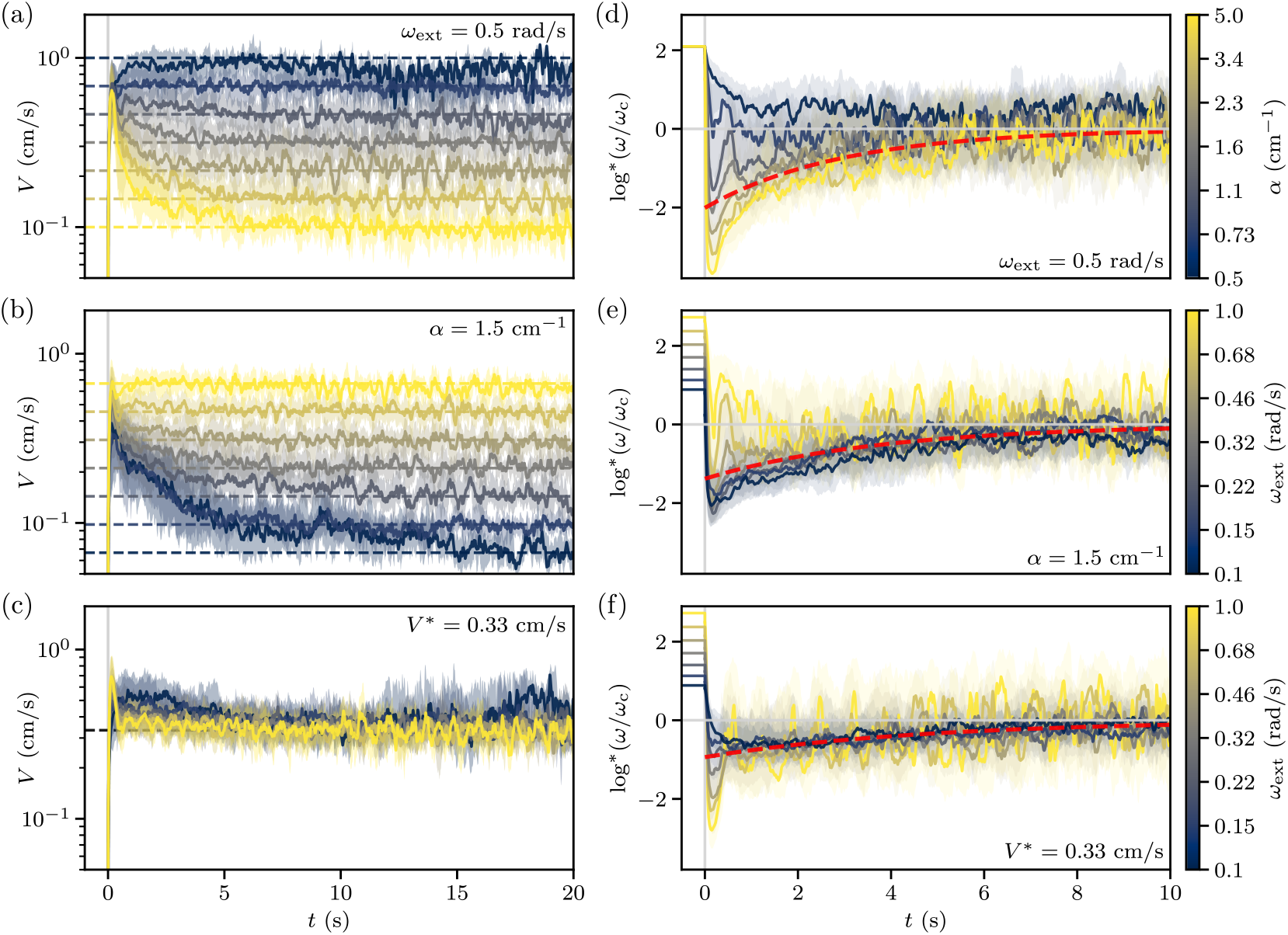
Initial speed transients in response to external currents for different values of feedback gain *α* and external flow rate *ω*_ext_, from the same experiments as in Fig. 2. (a) Traces of swimming speed *V* over time, aligned such that the initiation of swimming corresponds to *t* = 0, for the experiments with different values of *α* (color-coded values) with a constant *ω*_ext_. The swimming speed (solid lines and shaded areas indicate the quartiles of the distributions) converge to the corresponding target speeds (dashed lines). (b) Same as (a), but for the experiments with different values of *ω*_ext_ with a constant *α*. (c) Same as (a), but for experiments with different values of *α* and *ω*_ext_ such that their ratio is fixed. (d) Traces of sensory drive *S* = log^*^(*ω/ω*_c_) estimated from the speed transients of (a).The sensory drive (solid lines and shaded areas indicate the quartiles of the distributions) converges to zero approximately following an exponential decay (red dashed line, best fit). (e) Same as (d), but estimated from the speed transients of (b). (f) Same as (d), but estimated from the speed transients of (c).

If the goal of this behavioral reflex is to maintain the position of the fish stationary, this observation can be understood as follows. Before initiation of swimming, the fish perceives a sustained forward flow rate for a time *T*_r_, corresponding to a backward drift of its position by a distance *ω*_ext_*T*_r_*/α*. To correct for this backward drift, the fish must temporarily swim faster than the target speed in order to restore its position to the value prior to the appearance of the current. While Eq. 2 does not account for this phenomenon, we can propose a simple mechanism that does. We can introduce a second term that contributes to the evolution of the motor drive *M* while the fish is swimming. This correction term can simply be proportional to the accumulated sensory drive Φ, leading to the following equation:

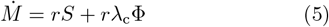

Where the rate *λ*_c_ determines the magnitude of the correction. This way, any kind of positional drift leads to an accumulation of Φ, which drives the motor drive to compensate for the drift while simultaneously driving Φ back to zero. The advantage of this mechanism is that it not only corrects for the drift accumulated before swimming initiation, but also compensates for other possible sources of drift, such as asymmetries in the response function, which have been observed experimentally in some fish [35].

If *λ*_c_ is small compared to the convergence rate of Eq. 3, Eq. 5 can be approximately solved using a separation of time scales. Over short time scales, we consider the second term in Eq. 5 to be effectively constant, while the dynamics of the first term lead to a quasi–steadystate. Considering 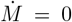, we find the quasi-steadystate sensory drive *S*_qss_ = − *λ*_c_Φ. Therefore, *V* will not converge toward *V* ^*^, but rather toward the quasi-steadystate fixed point:

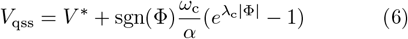

If the accumulated sensory drive is positive Φ > 0, then *V* converges to a value larger than *V* ^*^ in order to compensate for the accumulated drift. Conversely, if Φ < 0, meaning that the fish has been swimming faster than the current, then *V*_qss_ < *V* ^*^. Thus, over short time scales, *V* either converges toward *V*_qss_ or oscillates around it in a delay-induced limit cycle. To determine how this quasi–steady-state evolves over longer time scales, we consider the defining equation for the accumulated sensory drive 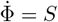, and substitute our result for *S*_qss_, finding 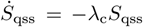. We therefore find that the quasisteady-state sensory drive decays exponentially to zero with a rate *λ*_c_.

Before swimming begins, Φ accumulates a positive sensory drive up to an average value of Φ_r_ (see Sec. III D), corresponding to a negative quasi-steady-state sensory drive at initiation *S*_qss_(0) = − *λ*_c_Φ_r_, and therefore *V*_qss_(0) > *V* ^*^. Afterwards, we expect the sensory drive to decay exponentially as 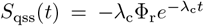. Indeed, we find that, after a quick transient following swimming initiation, the traces of *S* for most of the parameter values converge to a similar value, which then slowly approaches zero (Fig. 8d-f). This behavior is not observed for parameter values where the target speed *V* ^*^ is close to *V*_max_, as the fish cannot attain speeds significantly higher than the target. Excluding the experiments for which *V* ^*^ > 0.6 cm/s, we fitted the average sensory drive following swimming initiation with exponential decays. We found similar best fit values of Φ_r_ ≈ 5 s and *λ*_c_ ≈ 0.27 s^−1^ for the three different experimental protocols. The value of Φ_r_ agrees with the one obtained in Sec. III D when analyzing reaction times, while the value of *λ*_c_ provides a posteriori confirmation of the validity of our time scale separation hypothesis.

For simplicity, we have assumed a perfect integration Φ of the sensory drive *S*, but it could be reasonable to include a leak term, resulting in a leaky integration over a time scale of several seconds. This modification does not change the analysis qualitatively, but only slightly increases the decay rate of the quasi–steady-state solution.

## IV. STABILIZATION THROUGH INTERMITTENT LOCOMOTION

### A. Intermittent swimming in zebrafish

Here, we extend the framework that we developed for continuous locomotion in the previous section to the case of intermittent locomotion. Because of the universality of the principles underlying the nonlinear sensorimotor transformations illustrated in Fig. 5, we expect a similar control process to remain valid in the case of intermittent swimming. In this way, the same mechanism could underlie adaptive stabilization as danionella grow larger and transition to intermittent swimming [45], as well as in other species such as zebrafish. In fact, we will show that this framework can explain the experimental observations of visuomotor stabilization in the larval zebrafish (see Sec. II B). In this case, we hypothesize that the control process functions similarly to the case of continuous swimming (Fig. 5), in which the optic flow is first compressed and then integrated in time, accumulating into a motor drive *M*. The key difference is that, while in the continuous case *M* is directly reflected in the swimming speed *V*, in the intermittent case, *M* regulates the swimming speed *V* only at the times when the animal swims.

In freely swimming conditions, the swimming bouts of zebrafish exhibit a stereotyped speed profile, characterized by an initial peak lasting approximately ≈ 200 ms, followed by a decay to zero [24]. Upon normalizing each profile by its average speed *V*_b_, all traces collapse onto the same curve. Thus, if the bout duration *T*_b_ is defined using a speed threshold, one finds that *T*_b_ increases with the bout speed *V*_b_, whether the threshold is applied to the speed of the fish or of its tail movements [24, 38]. Because of the peaked shape of the speed profile the bout duration is not strongly modulated with bout speed, remaining of the order of *T*_b_ ≈ 200 ms, which we take as the nominal bout duration in the following. This observation suggests that the observed variations in *T*_b_ arise from corresponding variations in *V*_b_. Since the relative variations of *T*_b_ remain small compared to those of *V*_b_ we focus on the dynamics of the speed and assume a constant bout duration for simplicity.

It was shown in head-restrained experiments that the fish can adjust the speed of the tail mid-bout in response to perturbations of the visual feedback [23], but this modulation only takes place in the final part of the bout, therefore we do not expect it to significantly affect the speed of the fish. Here, we do not go into the details of how such a speed profile arises. Instead, we assume that a mechanism exists that terminates swimming shortly after it started, resulting in discrete swimming events. Nevertheless, it is worth noting that sensorimotor delays involved in the control of swimming speed have been proposed to play a role in determining such speed profiles and the termination of swimming underlying intermittent locomotion [45].

For simplicity, we consider the swimming speed to have a constant value *V*_b_ for the duration *T*_b_ of a bout (Fig. 9). Then, for an external current with a constant flow rate *ω*_ext_, the motor drive *M* will increase at a constant rate in the absence of visual feedback. When the fish swims, the net optic flow rate becomes *ω*_ext_ − *αV*_b_, decreasing the rate of change of *M*, which becomes negative for a backward optic flow. When the fish initiates a bout, the swimming speed *V*_b_ is determined by the value of the motor drive *M* at that instant.

**FIG. 9.**
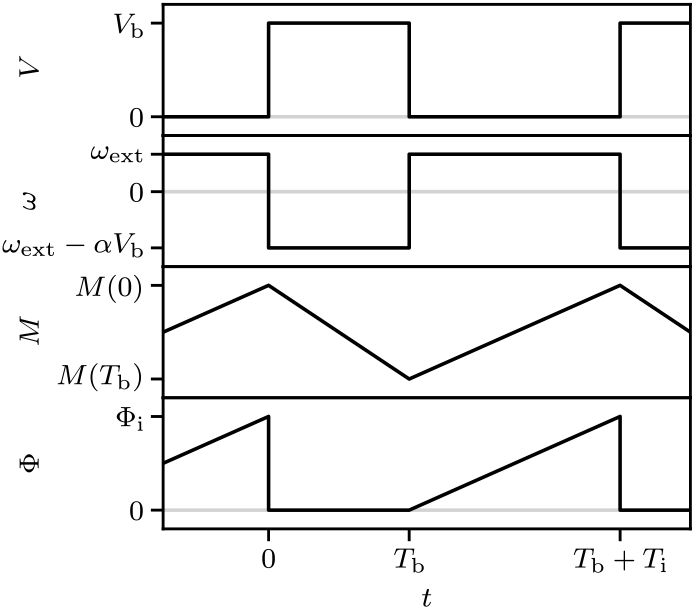
Schematic that illustrates the time dependent variables in a simplified model of intermittent swimming. The swimming speed *V* changes between 0 when the fish is not swimming for the duration *T*_i_ of the interbout interval to the bout speed *V*_b_ for the duration *T*_b_ of the bout. The motor drive *M* grows when the net optic flow rate *ω* is positive because of the external current and decreases when *ω* is negative because of the contribution of visual feedback. During the interbout interval, the accumulated sensory drive Φ increases linearly up to a threshold Φ_i_.

We do not consider the effect of sensorimotor delays in this case, as they do not significantly change the behavior of the system. Upon bout initiation, a short delay occurs before the tail movements affect the perceived optic flow and, consequently, the update of the motor drive. As a result, *M* continues to increase for a duration *τ* following the release of the motor command, after which the perceived visual feedback drives a different evolution for a duration *T*_b_. Thus, the sensorimotor delay only affects the interbout dynamics by introducing a refractory period during which the fish still perceives a delayed back-ward optic flow, even if it has already stopped swimming. Now, if the fish swims at speed *V*_*n*_ at the *n*-th bout, then we can simply compute the variation of *M* due to the net optic flow until the following bout and express the speed of the following bout in terms of that of the previous one *V*_*n*+1_ = *f* (*V*_*n*_). This gives us a discrete dynamical system for the evolution of the bout speed *V*_b_. We will investigate the behavior of the system for different forms of the function *f*, corresponding to either linear or nonlinear sensorimotor transformations. First, we look at the dynamics of bout initiation, which determine the interbout duration *T*_i_.

### B. Interbout duration in zebrafish

In freely swimming experiments where the feedback gain *α* was manipulated by changing the distance of the visual pattern, it was shown that the rate of swimming bouts *f*_b_ depends on the external flow rate *ω*_ext_, but not on *α* [24]. This suggests that the mechanism for bout initiation is independent of the intensity of visual feedback. While the bout speed *V*_b_ is governed by the motor drive *M*, whose value depends on *α*, the interbout duration *T*_i_ should only depend on *ω*_ext_.

Other experiments in restrained conditions have shown that *f*_b_ can also be modulated by changing *α* [21, 23]. This difference could be due to the fact that restraining the fish alters the observed response [38]. Here, we will follow the observations in freely swimming conditions and consider a mechanism for bout initiation that depends solely on *ω*_ext_.

We consider the simple accumulation model previously used for the initiation of swimming in danionella (Sec. III D). Here, we assume that the accumulated sensory drive Φ resets to a fixed value after each swimming bout, such that it only depends on the forward optic flow experienced during the interbout interval. Analogous to Eq. 4, the average interbout interval is then given by *T*_i_ = Φ_i_*/* log(*ω*_ext_*/ω*_th_) for *ω*_ext_ > *ω*_th_ (Fig. 9). Following the same reasoning as in Sec. III D, fluctuations in the sensory drive would lead to a lognormal distribution for *T*_i_. We used the same functional form as in Eq. 4 to fit the dependence of the interbout interval on the external flow rate from experiments on freely swimming zebrafish [38] (Fig. 10a). This fit captures the data with best-fit parameters Φ_i_ ≈ 1.1 s and *ω*_th_ ≈ 0.03 rad/s. The estimated value of *ω*_th_ is consistent with the one obtained for danionella in Sec. III D.

**FIG. 10.**
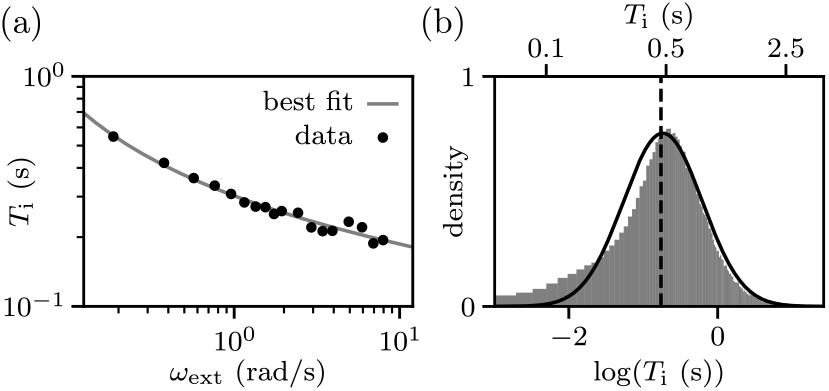
Interbout duration in larval zebrafish. (a) Interbout duration *T*_i_ in freely swimming larval zebrafish as a function of the flow rate *ω*_ext_ of an external visual current (black points). The data is well-fit by an inverse logarithmic dependence (gray line, see Eq. 4). Data from Ref. 38. (b) Histogram of the logarithm of the interbout durations *T*_i_ in freely swimming larval zebrafish in the absence of external visual currents. The dashed line indicates the median while the solid line is a Gaussian distribution which fits the data. Data from Ref. 46.

We can even extend this idea to spontaneous navigation in the absence of external currents, where larval zebrafish initiate swimming bouts with a variable interbout duration *T*_i_. The distribution of interbout durations is unimodal with a peak at ≈ 500 ms [46]. Experimentally measured histograms of *T*_i_ have been fitted with a negative binomial distribution. Although this fit captures the data, the negative binomial is intended to model a discrete random variable, whereas the interbout duration is continuous. Moreover, it provides no insight into the possible mechanisms underlying swimming initiation. We found that the same histogram can be fit equally well by a lognormal distribution, which is consistent with our hypothesized mechanism for swimming initiation (Fig. 10b). After a swimming bout terminates, an internal drive is integrated over time until it reaches a threshold sufficient to trigger the next bout. In the absence of salient external stimuli, we expect this internal drive to remain approximately constant, although it may be modulated over long time scales by external factors, such as temperature [47], or internal ones, such as hunger [48].

### C. Feedback control with linear transformations

Before examining the effects of the nonlinear transformations studied in the context of continuous locomotion (Fig. 5), we first analyze the behavior of the system in the absence of nonlinearities. We consider 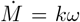, so that the motor drive *M* is simply proportional to the integral of the optic flow rate *ω* with responsiveness *k*, while the bout speed *V*_*n*_ = *M* (*t*_*n*_) directly reflects the motor drive at the time of the bout. As the fish swims with speed *V*_*n*_ at time *t*_*n*_, the motor drive changes first with rate *k*(*ω*_ext_ − *αV*_*n*_) for the duration of the visual feedback *T*_b_ and then with rate *kω*_ext_ for the duration of the interbout interval *T*_i_. We can then express the swimming speed of the following bout as:

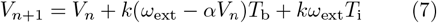

The fixed point of this linear difference equation can be found by setting *V*_*n*+1_ = *V*_*n*_, finding:

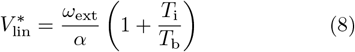

This is the speed at which the feedback flow rate exactly compensates the external flow rate over the interval between two consecutive bouts, corresponding to a fish that, on average, maintains a stable position. The solution to Eq. 7 can be written explicitly in terms of the initial condition *V*_0_ and the fixed point 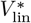 using the geometric sum formula [49]:

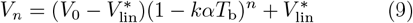

Eq. 9 holds for *kαT*_b_ ≠ 0, otherwise *V* simply grows by the same amount *kω*_ext_(*T*_b_ +*T*_i_) at each step. We see that the behavior of the solution depends on the dimensionless parameter combination *kαT*_b_. For 0 < *kαT*_b_ < 1 the speed converges exponentially to 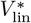. For 1 < *kαT*_b_ < 2 the speed converges to 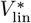 by alternating above and below it at each step. For *kαT*_b_ > 2 the fixed point becomes unstable and the speed diverges by alternating above and below it at each step.

To illustrate these three distinct dynamical regimes, we plot the solutions for three different values of *kαT*_b_ (Fig. 11). These solutions can be also constructed graphically as cobweb plots, by plotting *V*_*n*+1_ as a function of *V*_*n*_ and iterating the map by moving vertically from the identity line to this curve and horizontally back to the identity line to obtain successive bout speeds.

**FIG. 11.**
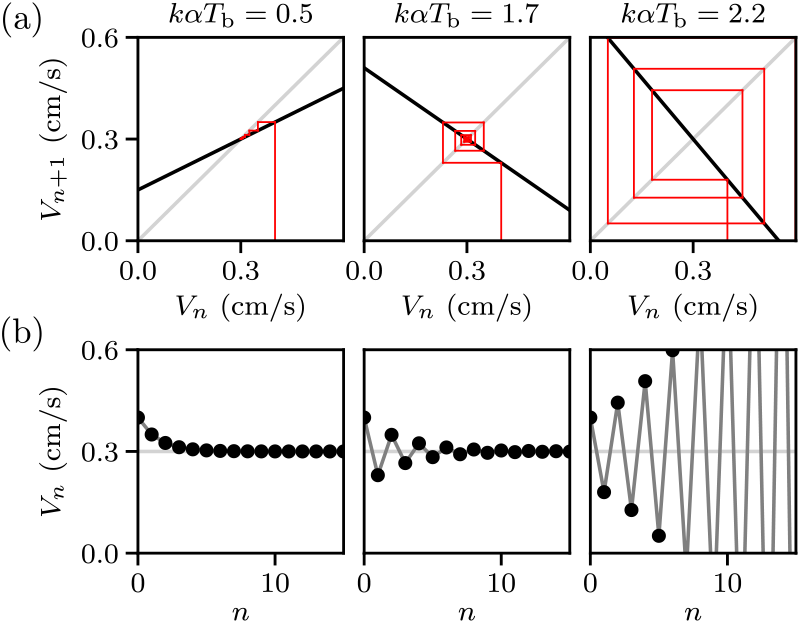
Examples of cobweb plots and swimming speed sequences for the linear difference equation. (a) Iterated maps (black lines) from Eq. 7 and corresponding cobweb constructions (red lines) for three different values of the parameter combination *kαT*_b_, with initial condition *V*_0_ = 0.4 cm/s and target speed 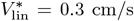. (b) Solutions of Eq. 7 corresponding to the cobweb construction in (a).

The parameter combination *kαT*_b_, which controls the system behavior, can be interpreted as the relative change in speed in the next bout caused by the visual feedback resulting from the current bout. If *kαT*_b_ > 1 we see that the system overcorrects and overshoots the target at each step, if *kαT*_b_ > 2 these overcorrections grow in size at each step, making the system unstable.

This instability is analogous to that discussed in the case of continuous locomotion (Sec. III B). The difference is that, in that case, the instability was due to the presence of sensorimotor delays, whereas here it results from fact that the fish can only control its speed at discrete times, due to the intrinsically intermittent nature of its locomotion. The system becomes unstable if either the responsiveness *k* or the feedback gain *α* become sufficiently large. While the parameter *k* is intrinsic to the fish, *α* depends on the specific environment surrounding the fish, indicating that same problem observed in Sec. III B also occurs in the case of intermittent locomotion. We now examine how this behavior changes when logarithmic transformations are considered, analogous to the analysis of Sec. III C.

### D. Feedback control with logarithmic transformations

We have seen that logarithmic transformations prevent dynamical instabilities in the case of continuous locomotion, here we investigate their effect on visuomotor stabilization during intermittent locomotion. We consider the same sensorimotor transformations as in Eq. 2, and derive a difference equation for the speed of a swimming bout in terms of that of the previous one, analogous to the linear case. Now, the motor drive changes with rate *r* log^*^((*ω*_ext_ − *αV*_*n*_)*/ω*_c_) for a time *T*_b_ and with rate *r* log^*^(*ω*_ext_*/ω*_c_) for a time *T*_i_, resulting in the following nonlinear difference equation for the swimming speed:

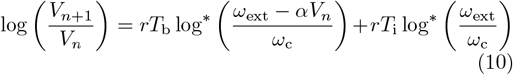

Eq. 10 can be further simplified using the fact that exp(log^*^(*x*)) = (1 + |*x*|)^sgn(*x*)^, leading to:

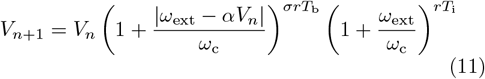

Where we defined *σ* = sgn(*ω*_ext_ − *αV*_*n*_). To stabilize its position, the fish must swim at a speed *V*_*n*_ larger than *ω*_ext_*/α*, corresponding to a backward net optic flow during the bout and *σ* = − 1.

Eq. 11 has a trivial fixed point 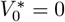, corresponding to the absence of swimming, and another one corresponding to a finite value of the speed 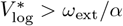:

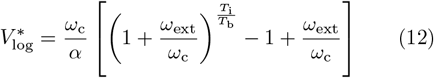

We note that this expression differs from the one corresponding to an exact stabilization of the position (Eq. 8), which was obtained as the fixed point 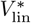 of the linear difference equation (Eq. 7). This difference is expected as in the nonlinear case the fish does not adjust its speed to maintain the optic flow rate *ω* constant on average, but rather its logarithmic representation *S*. The two solutions coincide only for *T*_i_ = *T*_b_, in which case the optic flow rates during swimming and resting periods are opposite, yielding 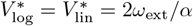.

To quantify the discrepancy, we define the ratio *η* between the fixed point of the nonlinear equation and that of the linear one (Fig. 12a):

**FIG. 12.**
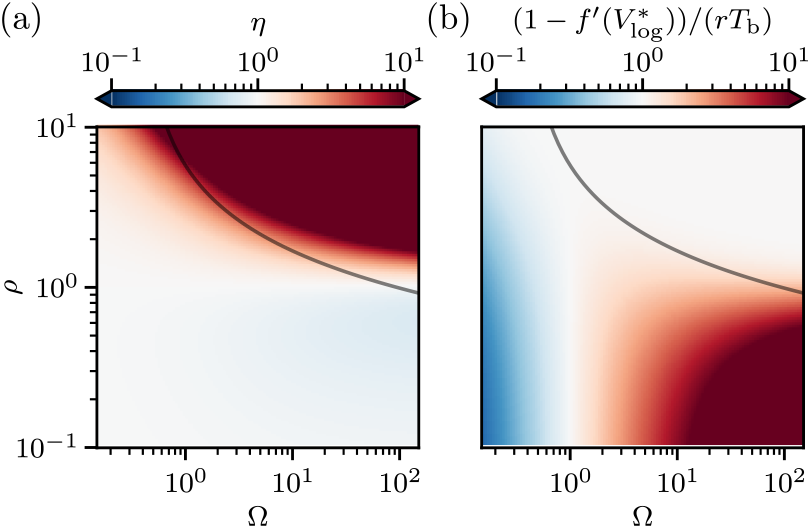
Parameter dependence of the fixed point of the nonlinear map (Eq. 11). (a) Ratio *η* between the fixed point of the nonlinear map and that of the linear one, as a function of the dimensionless parameter ratios *ρ* and Ω (Eq. 13). The relationship between *T*_i_ and *ω*_ext_ from the best fit of Fig. 10b is plotted as a solid line. (b) Deviation from the identity of the slope 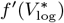 of the nonlinear map at the fixed point as a function of *ρ* and Ω (Eq. 15).

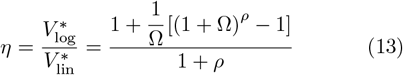

Which only depends on the ratios Ω = *ω*_ext_*/ω*_c_ and *ρ* = *T*_i_*/T*_b_. For *η* > 1, the fish swims faster than the target and moves forward on average, whereas for *η* < 1, it moves slower and drifts backward. We find that *η* = 1 when *ρ* = 1, meaning that the fish stabilizes its position exactly when *T*_i_ = *T*_b_. For *ρ* > 1 we find *η* > 1, with the fish swimming faster than the target and moving forward on average, whereas for *ρ* < 1 we find *η* < 1, with moving slower and drifting backward. In the limit Ω → 0, *η* → 1 independently of *ρ*. In the limit Ω → ∞, *η* → 1*/*(1 + *ρ*) for *ρ* < 1, whereas *η* → ∞ for *ρ* > 1.

Intuitively, the fish tends to swim at a speed that balances the logarithms of forward and backward optic flow rates, weighted by their corresponding durations. For large Ω, we can approximately write 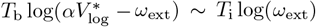, which leads to the scaling 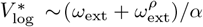, as in Eq. 12.

Because of its nonlinear nature, we cannot explicitly solve Eq. 11 as we did for Eq. 7. Nevertheless, we can study the stability of its fixed points to understand its behavior. To do so, the equation can be linearized in the vicinity of a fixed point, where it effectively behaves as a linear map Then, the behavior close to that fixed point depends on the slope of the map *V*_*n*+1_ = *f* (*V*_*n*_) (Eq. 11). If the absolute value of the slope is smaller than 1, the fixed point is stable and the solutions converge toward it. Otherwise, the fixed point is unstable and the solutions diverge away from it, either monotonically for a positive slope or alternating on each side for a negative one. Therefore, we have to study the slope of Eq. 11:

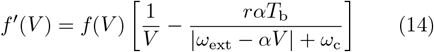

If we evaluate Eq. 14 at 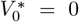, we find 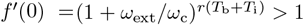, thus this fixed point is always unstable. The slope at the non-zero fixed point 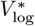 is instead given by (Fig. 12b):

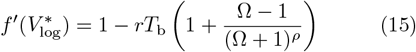

We thus find that the stability of this fixed point is independent of the feedback gain *α*. The fixed point can still become unstable when 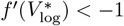, which can happen for large values of the system responsiveness *rT*_b_. The instability boundary depends on the values of Ω and *ρ*. For Ω → 0, 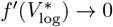 independently of *ρ*. For Ω → ∞, 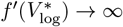 for *ρ* < 1 and 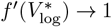 for *ρ* > 1.

We can also numerically simulate the system by choosing an initial condition *V*_0_ and iteratively evaluating Eq. 11 To illustrate the range of possible behaviors, we plotted solutions for different values of *rT*_b_, together with the corresponding cobweb constructions (Fig. 13). As the slope at the fixed point 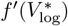 is increased, one first observes monotonic and then alternating convergence to 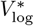. Eventually, the fixed point becomes unstable, however, as opposed to the linear system, the solutions do not diverge to infinity but instead exhibit either periodic trajectories or chaotic behavior.

**FIG. 13.**
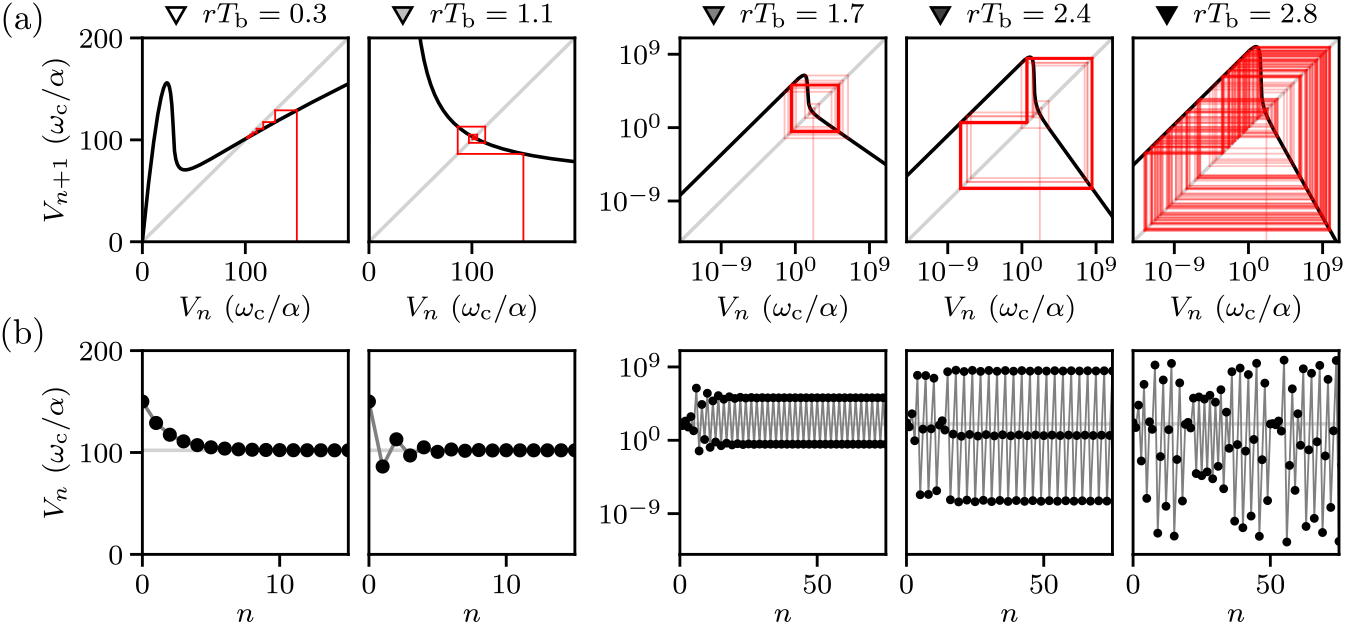
Examples of cobweb plots and swimming speed sequences for the nonlinear difference equation. (a) Iterated maps (black lines) from Eq. 11 and corresponding cobweb constructions (red lines) for five different values of the parameter combination *rT*_b_, with initial condition *V*_0_ = 150*ω*_c_*/α*, for Ω = 30, *ρ* = 1.25. (b) Solutions of Eq. 11 corresponding to the cobweb construction in (a). The triangles mark the corresponding values in the bifurcation diagram of Fig. 14.

To investigate the existence of periodic trajectories, we analyzed the behavior of the second iterate *f* ^2^(*V*), while varying the three independent parameters Ω, *ρ*, and *rT*_b_. In addition to the fixed points of *f* (*V*), we found new fixed points that appear when *rT*_b_ becomes sufficiently large. These points correspond to period-2 cycles, that is, oscillatory trajectories in which the swimming speed alternates between two different values. Such solutions can appear even before 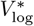 becomes unstable, resulting in a situation where multiple attractors coexist and the long term behavior of the solutions depends on the initial condition. Further increasing *rT*_b_, we also found additional fixed points of the third iterate *f* ^3^(*V*), corresponding to period-3 cycles. The existence of a period-3 cycle then also implies the existence of cycles of any other period and of chaotic behavior [50]. We constructed a bifurcation diagram of the system by simulating it over a range of values of *rT*_b_ and plotting the swimming speeds that are visited by the solutions after an initial transient (Fig. 14). These points indicate the attractors of the system as a function of *rT*_b_, that is, the sets of values to which the solutions tend to. We see that, as one increases *rT*_b_, the attractors change from a single stable fixed point to two points, corresponding to a period-2 cycle. Then, we find different intervals either containing a finite number of curves, corresponding to cycles with different periods, or bands that are dense with points, corresponding to chaotic attractors.

**FIG. 14.**
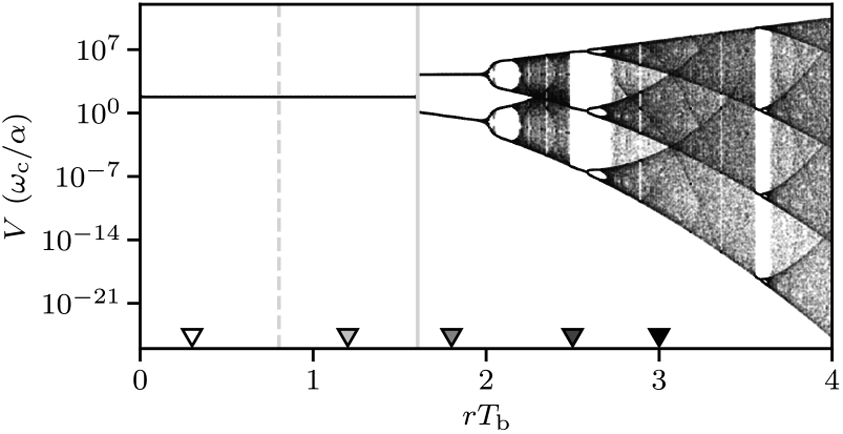
Bifurcation diagram showing the attractors of Eq. 11 as a function of *rT*_b_. It is obtained by plotting the points visited by the solutions of the equation after an initial transient. The triangles indicate the values of *rT*_b_ used in the examples of Fig. 13, the other parameters are the same as in that figure.

Now that we analyzed the behavior of the model, we can study whether it reproduces the experimental observations on the optomotor response in zebrafish larvae.

### E. Interpretation of experiments in zebrafish

#### 1. Dynamics of adaptation

Here we show that the adaptation to changes in feedback gain observed in Ref. 21 can be reproduced by considering the evolution of the bout speed *V*_b_ according to Eq. 11 (Fig. 15). By considering the same responsiveness rate *r* ≈ 1.6 s^−1^ that was estimated for danionella, we get *rT*_b_ ≈ 0.3. The other two parameters, *ρ* and Ω, are linked together by the interbout dynamics through *T*_i_ = Φ_i_*/* log(*ω*_ext_*/ω*_th_), as described in Sec. IV B (see curves in Fig. 12).

**FIG. 15.**
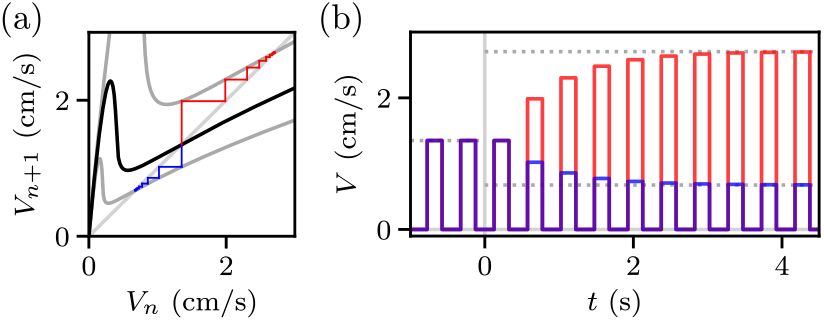
System response to changes of the feedback gain. (a) Cobweb constructions showing the evolution of the swimming speed according to Eq. 11 (black line) when the feedback gain is changed from *α*_0_ = 4 cm^−1^ to either *α*_+_ = 8 cm^−1^ (blue lines) or *α*_−_ = 2 cm^−1^ (red lines). Parameter values: *ω*_ext_ = 2 rad/s, *T*_b_ = 0.2 s, *T*_i_ = 0.25 s. (b) Swimming speed traces over time corresponding to the two solutions shown in (a), showing how the bout speed progressively changes over several bouts from the old fixed point to the new ones (dotted lines) after the feedback gain is changed (at *t* = 0).

Using these values, we find that for *ω*_ext_ in the range 0.1–10 rad/s the slope of *f* (*V*) at the fixed point 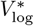 (Eq. 15) remains in the range 0.3–0.7, corresponding to a monotonic convergence to the fixed point (Fig. 13, left). Then, for fixed values of the parameters *α* and *ω*_ext_, the bout speed converges to the fixed point 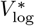 and the fish keeps swimming at that speed indefinitely. Upon changing the feedback gain *α*, the map *f* (*V*) is rescaled, resulting in a different equation for the dynamics of *V*_b_ and a new fixed point 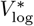. The swimming speed then evolves toward this new fixed point with a convergence rate determined by the slope 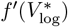, since in this regime the function *f* (*V*) can be reasonably approximated by its linearization around the fixed point. The bout speed converges exponentially to the new fixed point (Fig. 15), reaching 90% of the way after 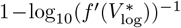 bouts.

If we consider *ω*_ext_ = 2 rad/s, as in the experiments of Ref. [21], we find that it converges after ≈ 5 bouts, which matches the time scale of the observed adaptation. Such a gradual change in swimming speed over several bouts has been interpreted as reflecting motor learning of the new feedback gain, but we find that it can also arise from the dynamics of a simple feedback controller.

We note that for linear control (Eq. 7), the convergence rate depends on the feedback gain, with the system becoming unstable if *α* is larger than a critical value. In contrast, the nonlinear transformation included in Eq. 10 make the system adaptive, such that the convergence rate is independent of *α*.

#### 2. Positional stabilization

In Sec. IV C, we have shown that a linear feedback controller (Eq. 7) leads to an instability for large values of the feedback gain *α*, something that was not addressed before by studies that proposed models of this kind [24]. Then, in Sec. IV D, we demonstrated that logarithmic transformations (Eq. 10) prevent this instability by making the stability of the system independent of *α*. Yet, while the solutions of the linear system (Eq. 8) corresponds to a perfect positional stabilization, the nonlinear system can converge to different values (Eq. 12), as quantified by the ratio *η* (Eq. 13).

To study the stabilization in the case of the nonlinear system (Eq. 10) we can simulate its behavior across different values of the external flow rate *ω*_ext_ and the feedback gain *α*. Up to this point, we considered deterministic dynamics for the swimming speed, however, the behavior of the fish is inherently stochastic. One part of this stochasticity can be recovered by considering the interbout duration *T*_i_ as a lognormally distributed random variable, rather than just using its mean value as we have done so far. Additionally, we can consider another source of noise in the evolution of the motor drive *M*, that until now we took to be deterministic. Similarly to Ref. 35, we can add a Gaussian white noise term in the rate of change of *M* with amplitude *σ*_*ξ*_. Integrating this stochastic term in the time from one bout to the next we obtain an additional noise term in Eq. 10 which is distributed as a Gaussian random variable with zero mean and standard deviation 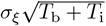.

Taking into account these two sources of noise, we can numerically simulate trajectories of the swimming speed starting with some initial condition *V*_0_ and then sampling the interbout duration *T*_i_ for each bout and the swimming speed *V*_n+1_ at the following bout. Such simulations were performed for different values of *ω*_ext_ and *α*, matching those used in the experiments of Refs. We considered the value of *σ*_*ξ*_ found in danionella from Ref. 35, whereas *σ*_i_ was estimated from the standard deviation of log(*T*_i_) in zebrafish from Refs. 20, 21 and 38.

To understand whether the solutions correspond to an actual stabilization of the fish position, we should not simply look at the bout speed *V*_b_, but rather at the average speed including swimming and resting periods ⟨*V*⟩ = *V*_b_*/*(1 + *ρ*) (Fig. 16, blue). We see that the average speed of the fish varies in accordance with the target speed *V* ^*^ = *ω*_ext_*/α*, but remains above it across the range of parameters considered. This is expected as we have seen in the previous section that for *T*_b_ ≠ *T*_i_ the solutions converge to a value of the speed 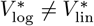 that does not result in a perfect stabilization of the fish. We see that the discrepancy becomes larger for smaller values of *ω*_ext_ (see Fig. 12a). We find that increasing *ω*_ext_ from 0.2 rad/s to 8 rad/s, the ratio *η* decreases from 4 to 1.

**FIG. 16.**
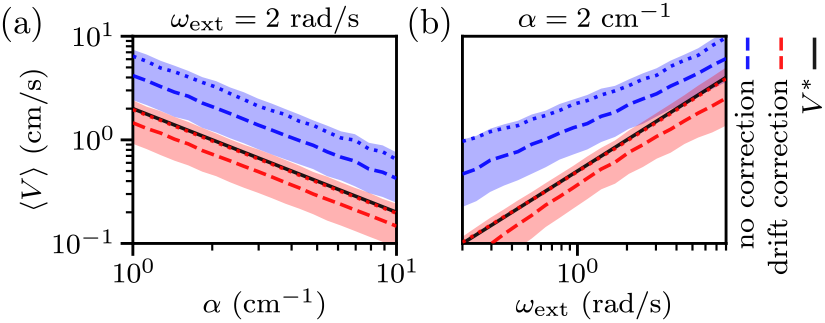
Average swimming speed ⟨*V*⟩ for simulations of Eq. 11 (dashed lines and shaded areas indicate the quartiles of the distributions over simulated bouts, dotted lines indicate the mean), compared with the target speed *V* ^*^ = *ω*_ext_*/α* (black solid lines). The relation between *T*_i_ and *ω*_ext_ is taken from the fit in Fig. 10b. Stochasticity is included both in the computation of interbout duration (*σ*_*i*_ = 0.3) and bout speed (*σ*_*ξ*_ = 0.9 s^−1*/*2^). Results are shown either with a linear drift correction term (red, *λ*_c_ = 0.01) or without it (blue). (a) For different values of the feedback gain *α* and *ω*_ext_ = 2 rad/s, as in Refs. 20 and 21. (b) For different values of the external flow rate *ω*_ext_ and *α* = 2 cm^−1^, as in Ref. 38.

While some experiments in zebrafish have reported an imperfect stabilization and a net drift in the position of the fish [24], which could be partly explained by the observed discrepancy, others report perfect stabilization [38], suggesting that another mechanism might be at play. Similarly to what we observed in Sec. III E, we could expect an additional mechanism that integrates the optic flow rate *ω* on a longer time scale and corrects an eventual positional drift. As the imperfect stabilization results from the compressed representation of the flow rate, the drift correction term would have to be proportional to the integral of *ω*, rather than to that of *S* as it was the case in in Sec. III E. Thus we added an additional term of the form *rλ*_c_ *∫ ω*(*t*)*dt* to the evolution of the motor drive and again simulated the dynamics (Fig. 16, red). We find that even a small correction term compensates for the discrepancy over time, resulting in an average swimming speed ⟨*V*⟩ that matches the target *V* ^*^ and therefore a perfect positional stabilization.

#### 3. Directional asymmetry in responsiveness

In Refs. 24 and 25, it was suggested that the fish only modulate their locomotion based on the forward optic flow corresponding to the external current, disregarding visual feedback. However, such an interpretation cannot explain how the fish can stabilize their position in different conditions, as the target speed does not only depend on the external flow rate *ω*_ext_, but also on the feedback gain *α*.

Here, we show how this interpretation can emerge when fitting a linear model of the form of Eq. 7 to data generated by the nonlinear dynamics of Eq. 11. Specifically, they considered a linear model with different responsiveness values, *k*_+_ and *k*_−_, for forward and backward flows, respectively. Fitting this model to the experimental data yielded large ratios of *k*_+_*/k*_−_, which were then interpreted as indicating that swimming speed is mainly determined by the forward flow.

To compare Eq. 11 with the linear model we can linearize the left hand side for small deviations from the fixed point 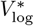. Then, the values of *k*_+_ and *k*_−_ can be read off by from the two terms in the update of the swimming speed, which are proportional to *T*_i_ and *T*_b_, respectively. We find 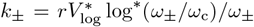, with *ω*_+_ = *ω*_ext_ and 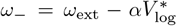. Finally, using the expression for the fixed point 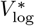 (Eq. 12), we get the following for the ratio between the two values:

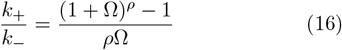

We see that *k*_+_*/k*_−_ > 1 for *ρ* > 1, whereas *k*_+_*/k*_−_ < 1 for *ρ* < 1. This is because for *ρ* > 1 the interbout lasts longer than the bout and thus the fish needs to swim at large speeds to compensate for the forward optic flow accumulated during the interbout interval, resulting in *ω*_−_ > *ω*_+_. As the rate of change of the motor drive depends on the logarithm of the optic flow rate, which is a sublinear function of *ω*, we find that the estimated linear responsiveness decreases with *ω*. Thus, a larger flow rate during the bout *ω*_−_ as compared to that during the interbout *ω*_+_, results in smaller backward responsiveness *k*_−_ compared to the forward one *k*_+_. Using the interbout durations corresponding to values of *ω*_ext_ in the range 0.1–1 rad/s, we obtain ratios *k*_+_*/k*_−_ in the range 3–7.

This shows that interpreting the dynamics of the nonlinear system through a linear model with asymmetric responsiveness can give the misleading impression that the fish is more sensitive to forward than backward flows, whereas the apparent asymmetry actually arises from the nonlinear compression of flow rate combined with the different durations of swimming and resting periods.

#### 4. Different integration time scales

Here, we show that the fact that bout initiation and bout speed are governed by distinct processes can account for the observations reported in Ref. 25 without the need to introduce a model with two separate integration time scales. The bout-triggered average stimulus that they obtained through reverse correlation can be explained by the integrate-to-threshold mechanism for bout initiation. This mechanism results in a first-passage problem for a random walk driven by the stochastically switching current direction, with a characteristic time scale that depends on the statistics of the stimulus.

They also showed that the swimming speed in response to closed-loop currents depends on the direction of a preceding open-loop current, even after a long pause. This second observation is consistent with our proposed mechanism for bout speed selection, which depends on the sensory drive integrated over time, without any leak term. We can actually expect a small leak term to be present, which we excluded here to keep the model as simple as possible, but which would account for the gradual decay of the effect of the open-loop stimulus as the pause duration increases.

Although consistent with the results of Ref. 22, our interpretation does not imply that the fish maintain a memory of their location in the environment, but only that they integrate the perceived optic flow rate over time, a mechanism compatible with the neural activity observed in that study.

## V. DISCUSSION

In this paper, we studied how the logarithmic encoding of physical variables in the brain can lead to robust positional stabilization through simple feedback control of motor output based on visual input. Such logarithmic encoding is a fundamental feature of nervous systems, widespread across the animal kingdom. At the sensory end it is known as the Weber-Fechner law [36], while at the motor end it corresponds to Hennemann’s size principle [37]. Although these two principles have often been discussed in terms of informational efficiency [51–53], here we have shown how they jointly contribute to sensorimotor computations. Our results demonstrate that these nonlinear sensorimotor transformations naturally result in an adaptive responsiveness to different contexts, that is, different relationships between motor output and reafferent visual feedback. This emergent adaptation prevents the dynamical instabilities that can arise because the response of any animal is inherently constrained by temporal delays between perception and action. These delays arise from finite processing times within the sensorimotor loop and, in many species, from the intermittent or cyclic nature of locomotion.

These ideas were first introduced in Ref. 35, which studied the optomotor response in danionella larvae, where the continuous modulation of swimming made it straightforward to infer the underlying sensorimotor transformations. Although visuomotor stabilization has been extensively studied in larval zebrafish, the underlying mechanisms have remained elusive due to the intermittent nature of their swimming. Behavioral readouts are available only during discrete swimming bouts, obscuring the continuous processes that contribute to stabilization. Here, we extended the concepts originally developed for danionella to formulate a mathematical framework for visuomotor stabilization, applicable to both continuous and intermittent locomotion.

In the continuous case, we obtained a delay differential equation describing the evolution of the speed and identified a potential instability arising from the delays in the sensorimotor loop. We showed how logarithmic transformations prevent this instability by making the dynamics multiplicative and limiting the responsiveness of the system, such that the stability becomes independent of the feedback gain and the solutions remain confined to limit cycles. We further analyzed the experimental data in danionella, modeling the initiation of swimming as an accumulation process that integrates the sensory drive up to a threshold. Finally, we identified an integration mechanism that compensates for positional drift.

In the intermittent case, we derived a difference equation for the speed and identified an instability analogous to the continuous case, but arising from the delays in between discrete locomotor events. Again, logarithmic transformations make the stability of the system independent of the feedback gain, depending only on parameters intrinsic to the animal and not on the environmental context. We used this framework to interpret the experimental evidence from the visuomotor literature on zebrafish larvae. Our model reproduced many experimental observations while remaining simple and using only a few parameters. Remarkably, using the same response function estimated for danionella, we still obtained results that closely match the observations in zebrafish.

Both danionella and zebrafish larvae allow for wholebrain functional imaging at cellular resolution and are used as model organisms in systems neuroscience to study how sensorimotor computations are implemented at the level of the neural activity [26, 54]. While this approach offers insight into the neural circuits underlying these behaviors, the high dimensionality of the recorded data and the limited temporal resolution of current imaging techniques make it challenging to directly infer the sensorimotor transformations [16, 55, 56]. This is evident from the fact that, although many of the studies described in Sec. II B recorded neural activity alongside visuomotor behavior in larval zebrafish, they still reached different conclusions regarding the mechanisms underlying stabilization [21–23, 25, 38].

While we applied this theory to explain visuomotor stabilization in two fish species, we expect it to be readily applicable to other animals. This is because such visuomotor reflexes [11, 57, 58], along with the principles of logarithmic coding [59], are found across many different species and contexts. In engineering, designing control mechanisms that rely solely on visual input remains notoriously difficult, and additional sensors are typically used to disentangle speed and distance from the measured optic flow [10, 60]. Our model, however, could be implemented as a simple algorithm that first compresses logarithmically the inputs and then decompresses them to generate the output. We suggest that this approach could enable robust stabilization in flying or swimming robots subjected to unpredictable external perturbations.

## ACKNOWLEDGMENTS

I thank the members of the Laboratoire Jean Perrin and the SmartNets European Training Network, especially Georges Debrégeas and Monica Coraggioso, for helpful feedback and discussions. This work was supported by the European Union’s Horizon 2020 research and innovation programme under the Marie Skłodowska-Curie grant agreement No. 860949 and by the Agence Nationale de la Recherche under the Locomat project ANR-21-CE16-0037.

## DATA AVAILABILITY

The analyzed data from experiments on danionella was previously published and is available on Zenodo [61]. The code written for the analysis has been deposited in a public repository on Zenodo [62].

## References

[1] M. F. Land and D.-E. Nilsson, Animal Eyes, second edition ed., Oxford animal biology series (Oxford University Press, Oxford, 2012).

[2] J. J. Gibson, Visually Controlled Locomotion and Visual Orientation in Animals, British Journal of Psychology 49, 182 (1958).

[3] R. Shadmehr, M. A. Smith, and J. W. Krakauer, Error Correction, Sensory Prediction, and Adaptation in Motor Control, Annual Review of Neuroscience 33, 89 (2010).

[4] S. G. Tzafestas, Mobile Robot Control and Navigation: A Global Overview, Journal of Intelligent & Robotic Systems 91, 35 (2018).

[5] R. C. Miall and D. M. Wolpert, Forward Models for Physiological Motor Control, Neural Networks Four Major Hypotheses in Neuroscience, 9, 1265 (1996).

[6] M. Kawato, Internal models for motor control and trajectory planning, Current Opinion in Neurobiology 9, 718 (1999).

[7] C. Tin and C.-S. Poon, Internal models in sensorimotor integration: perspectives from adaptive control theory, Journal of Neural Engineering 2, S147 (2005).

[8] D. Nguyen-Tuong and J. Peters, Model learning for robot control: a survey, Cognitive Processing 12, 319 (2011).

[9] J. J. Koenderink, Optic flow, Vision Research 26, 161 (1986).

[10] J. R. Serres and F. Ruffier, Optic flow-based collision-free strategies: From insects to robots, Arthropod Structure & Development From Insects to Robots, 46, 703 (2017).

[11] M. V. Srinivasan and S. Zhang, Visual motor computations in insects, Annual Review of Neuroscience 27, 679 (2004).

[12] G. P. Arnold, Rheotropism in fishes, Biological Reviews 49, 515 (1974).

[13] G. D. McCann, G. F. MacGinitie, and J. W. S. Pringle, Optomotor response studies of insect vision, Proceedings of the Royal Society of London. Series B. Biological Sciences 163, 369 (1997), publisher: Royal Society.

[14] E. Shaw and A. Tucker, The optomotor reaction of schooling carangid fishes, Animal Behaviour 13, 330 (1965).

[15] R. Portugues and F. Engert, The neural basis of visual behaviors in the larval zebrafish, Current Opinion in Neurobiology 19, 644 (2009).

[16] M. B. Orger and G. G. d. Polavieja, Zebrafish Behavior: Opportunities and Challenges, Annual Review of Neuroscience 40, 125 (2017), publisher: Annual Reviews.

[17] J. H. Bollmann, The Zebrafish Visual System: From Circuits to Behavior, Annual Review of Vision Science 5, 269 (2019).

[18] M. B. Ahrens, M. B. Orger, D. N. Robson, J. M. Li, and P. J. Keller, Whole-brain functional imaging at cellular resolution using light-sheet microscopy, Nature Methods 10, 413 (2013).

[19] T. Panier, S. A. Romano, R. Olive, T. Pietri, G. Sumbre, R. Candelier, and G. Debrégeas, Fast functional imaging of multiple brain regions in intact zebrafish larvae using Selective Plane Illumination Microscopy, Frontiers in Neural Circuits 7, 10.3389/fncir.2013.00065 (2013).

[20] R. Portugues and F. Engert, Adaptive Locomotor Behavior in Larval Zebrafish, Frontiers in Systems Neuroscience 5, 10.3389/fnsys.2011.00072 (2011).

[21] M. B. Ahrens, J. M. Li, M. B. Orger, D. N. Robson, A. F. Schier, F. Engert, and R. Portugues, Brain-wide neuronal dynamics during motor adaptation in zebrafish, Nature 485, 471 (2012).

[22] E. Yang, M. F. Zwart, B. James, M. Rubinov, Z. Wei, S. Narayan, N. Vladimirov, B. D. Mensh, J. E. Fitzgerald, and M. B. Ahrens, A brainstem integrator for self-location memory and positional homeostasis in zebrafish, Cell 185, 5011 (2022).

[23] D. A. Markov, L. Petrucco, A. M. Kist, and R. Portugues, A cerebellar internal model calibrates a feedback controller involved in sensorimotor control, Nature Communications 12, 6694 (2021).

[24] J. G. Holman, W. W. K. Lai, P. Pichler, D. Saska, L. Lagnado, and C. L. Buckley, A behavioral and modeling study of control algorithms underlying the translational optomotor response in larval zebrafish with implications for neural circuit function, PLOS Computational Biology 19, e1010924 (2023).

[25] R. Tanaka and R. Portugues, Algorithmic dissection of optic flow memory in larval zebrafish, Current Biology 35, 4870 (2025).

[26] L. Schulze, J. Henninger, M. Kadobianskyi, T. Chaigne, A. I. Faustino, N. Hakiy, S. Albadri, M. Schuelke, L. Maler, F. Del Bene, and B. Judkewitz, Transparent Danionella translucida as a genetically tractable vertebrate brain model, Nature Methods 15, 977 (2018).

[27] R. Britz, K. W. Conway, and L. Rüber, The emerging vertebrate model species for neurophysiological studies is Danionella cerebrum, new species (Teleostei: Cyprinidae), Scientific Reports 11, 18942 (2021).

[28] M. Hoffmann, J. Henninger, J. Veith, L. Richter, and B. Judkewitz, Blazed oblique plane microscopy reveals scale-invariant inference of brain-wide population activity, Nature Communications 14, 8019 (2023), number: 1 Publisher: Nature Publishing Group.

[29] T. J. Lee and K. L. Briggman, Visually guided and context-dependent spatial navigation in the translucent fish Danionella cerebrum, Current Biology 33, 5467 (2023).

[30] V. A. N. O. Cook, A. H. Groneberg, M. Hoffmann, M. Kadobianskyi, J. Veith, L. Schulze, J. Henninger, R. Britz, and B. Judkewitz, Ultrafast sound production mechanism in one of the smallest vertebrates, Proceedings of the National Academy of Sciences 121, e2314017121 (2024), publisher: Proceedings of the National Academy of Sciences.

[31] J. Veith, T. Chaigne, A. Svanidze, L. E. Dressler, M. Hoffmann, B. Gerhardt, and B. Judkewitz, The mechanism for directional hearing in fish, Nature 631, 118 (2024).

[32] D. Zada, L. Schulze, J.-H. Yu, P. Tarabishi, J. L. Napoli, J. Milan, and M. Lovett-Barron, Development of neural circuits for social motion perception in schooling fish, Current Biology 34, 3380 (2024), publisher: Elsevier.

[33] A. H. Groneberg, L. E. Dressler, M. Kadobianskyi, J. Müller, and B. Judkewitz, Development of sound production in Danionella cerebrum, Journal of Experimental Biology 227, jeb247782 (2024).

[34] G. Rajan, J. Lafaye, G. Faini, M. Carbo-Tano, K. Duroure, D. Tanese, T. Panier, R. Candelier, J. Henninger, R. Britz, B. Judkewitz, C. Gebhardt, V. Emiliani, G. Debregeas, C. Wyart, and F. Del Bene, Evolutionary divergence of locomotion in two related vertebrate species, Cell Reports 38, 110585 (2022).

[35] L. Demarchi, M. Coraggioso, A. Hubert, T. Panier, G. Morvan-Dubois, V. Bormuth, and G. Debrégeas, Logarithmic coding leads to adaptive stabilization in the presence of sensorimotor delays, Proceedings of the National Academy of Sciences 122, e2510385122 (2025), publisher: Proceedings of the National Academy of Sciences.

[36] J. C. Baird and E. J. Noma, Fundamentals of Scaling and Psychophysics (Wiley, 1978).

[37] E. Henneman, Relation between Size of Neurons and Their Susceptibility to Discharge, Science 126, 1345 (1957), publisher: American Association for the Advancement of Science.

[38] K. E. Severi, R. Portugues, J. C. Marques, D. M. O’Malley, M. B. Orger, and F. Engert, Neural Control and Modulation of Swimming Speed in the Larval Zebrafish, Neuron 83, 692 (2014).

[39] R. Portugues, M. Haesemeyer, M. L. Blum, and F. Engert, Whole-field visual motion drives swimming in larval zebrafish via a stochastic process, Journal of Experimental Biology 218, 1433 (2015).

[40] T. W. Dunn and J. E. Fitzgerald, Correcting for physical distortions in visual stimuli improves reproducibility in zebrafish neuroscience, eLife 9, e53684 (2020).

[41] S. D. Brown and A. Heathcote, The simplest complete model of choice response time: Linear ballistic accumulation, Cognitive Psychology 57, 153 (2008).

[42] C. Donkin and L. Van Maanen, Piéron’s Law is not just an artifact of the response mechanism, Journal of Mathematical Psychology 62-63, 22 (2014).

[43] G. Buzsáki and K. Mizuseki, The log-dynamic brain: how skewed distributions affect network operations, Nature Reviews Neuroscience 15, 264 (2014).

[44] R. Ulrich and J. Miller, Information processing models generating lognormally distributed reaction times, Journal of Mathematical Psychology 37, 513 (1993), place: Netherlands Publisher: Elsevier Science.

[45] M. Coraggioso, L. Demarchi, R. Wong, V. Dichio, C. Chaumeton, T. Panier, G. Morvan-Dubois, G. Goodhill, V. Bormuth, and G. Debrégeas, A sensorimotor instability drives a locomotor transition during fish development (2025), iSSN: 2692-8205 Pages: 2025.10.10.681567 Section: New Results.

[46] R. E. Johnson, S. Linderman, T. Panier, C. L. Wee, E. Song, K. J. Herrera, A. Miller, and F. Engert, Probabilistic Models of Larval Zebrafish Behavior Reveal Structure on Many Scales, Current Biology 30, 70 (2020).

[47] G. L. Goc, J. Lafaye, S. Karpenko, V. Bormuth, R. Candelier, and G. Debrégeas, Thermal modulation of Zebrafish exploratory statistics reveals constraints on individual behavioral variability, BMC Biology 19, 208 (2021).

[48] D. Clift, H. Richendrfer, R. J. Thorn, R. M. Colwill, and R. Creton, High-Throughput Analysis of Behavior in Zebrafish Larvae: Effects of Feeding, Zebrafish 11, 455 (2014), publisher: Mary Ann Liebert, Inc., publishers.

[49] S. Elaydi, An introduction to difference equations, 3rd ed., Undergraduate texts in mathematics (Springer, New York, 2005).

[50] T.-Y. Li and J. A. Yorke, Period Three Implies Chaos, The American Mathematical Monthly 82, 985 (1975), publisher: Taylor & Francis eprint: 10.1080/00029890.1975.11994008.

[51] H. Hatze, A teleological explanation of Weber’s law and the motor unit size law, Bulletin of Mathematical Biology 41, 407 (1979).

[52] W. Senn, K. Wyler, H. P. Clamann, J. Kleinle, H. R. Lüscher, and L. Müller, Size principle and information theory, Biological Cybernetics 76, 11 (1997).

[53] R. D. Portugal and B. F. Svaiter, Weber-Fechner Law and the Optimality of the Logarithmic Scale, Minds and Machines 21, 73 (2011).

[54] R. W. Friedrich, G. A. Jacobson, and P. Zhu, Circuit Neuroscience in Zebrafish, Current Biology 20, R371 (2010).

[55] T. Rose, P. M. Goltstein, R. Portugues, and O. Griesbeck, Putting a finishing touch on GECIs, Frontiers in Molecular Neuroscience 7, 10.3389/fnmol.2014.00088 (2014).

[56] F. Ali and A. C. Kwan, Interpreting in vivo calcium signals from neuronal cell bodies, axons, and dendrites: a review, Neurophotonics 7, 1 (2019).

[57] F. R. Harden Jones, The Reaction of Fish to Moving Backgrounds, Journal of Experimental Biology 40, 437 (1963).

[58] A. S. Mauss and A. Borst, Optic flow-based course control in insects, Current Opinion in Neurobiology 60, 21 (2020).

[59] E. R. Kandel, J. Koester, S. Mack, and S. Siegelbaum, eds., Principles of neural science, sixth edition ed. (McGraw Hill, New York, 2021).

[60] H. Chao, Y. Gu, and M. Napolitano, A Survey of Optical Flow Techniques for Robotics Navigation Applications, Journal of Intelligent & Robotic Systems 73, 361 (2014).

[61] L. Demarchi, M. Coraggioso, A. Hubert, T. Panier, G. Morvan-Dubois, V. Bormuth, and G. Debrégeas, Data and code - Logarithmic coding leads to adaptive stabilization in the presence of sensorimotor delays (2025).

[62] L. Demarchi, Code - A logarithmic theory of visuomotor stabilization (2025).

